# MISO: Microfluidic protein isolation enables single particle cryo-EM structure determination from a single cell colony

**DOI:** 10.1101/2025.01.10.632437

**Authors:** Gangadhar Eluru, Steven De Gieter, Stephan Schenck, Annelore Stroobants, Binesh Shrestha, Paul Erbel, Janine D. Brunner, Rouslan G. Efremov

## Abstract

Single particle cryo-EM enables reconstructing near-atomic or even atomic resolution 3D maps of proteins by visualizing thousands to a few million purified protein particles embedded in nanometer-thick vitreous ice. This corresponds to picograms of purified protein, which can potentially be isolated from a few thousand cells. Hence, cryo-EM holds the potential of one of the most sensitive analytical methods that deliver a high-resolution protein structure as a readout. In practice, more than a million times more starting biological material is required to prepare cryo-EM grids. To close the gap, we developed a micro isolation (MISO) method that combines microfluidics-based protein purification with cryo-EM grid preparation. We validated the method on soluble bacterial and eukaryotic membrane proteins. We showed that MISO enables protein structure determination starting from below one microgram of a target protein and going from cells to cryo-EM grids within a few hours. This scales down the purification by a factor of a few hundred to a few thousand and opens possibilities for the structural characterization of hitherto inaccessible proteins.

## Introduction

Electron cryogenic microscopy (cryo-EM) has become the mainstream technique to determine experimental structures of biological macromolecular complexes, including membrane proteins, fragile and flexible multicomponent complexes as well as protein complexes with pharmacologically relevant small molecules [1,2]. Theoretically, only around 10,000 randomly oriented particles are sufficient to reconstruct a protein map to atomic resolution [3]. In practice, the structures are solved after imaging a few hundred thousand to a few million protein particles [4]. Even then, this number of particles corresponds to only picogram amounts of imaged protein. Hence, single particle cryo-EM holds the potential to become a very sensitive analytical technique that delivers 3D atomic structures of the analytes. However, this requires that picogram amounts of proteins are purified to homogeneity and applied directly to a grid, which is currently not possible.

Recent methodological developments in cryo-EM led to the emergence of more stable and automated microscopes, efficient electron detectors [5], methods for automated data collection [6–9] image processing [10–12], protein model building and structure refinement [13–16]. These advances increased the throughput of electron microscopy workflow from imaging to refined 3D models. Furthermore, cheaper 100 kV high-resolution electron microscopes [17,18] hold promise to further improve the efficiency of imaging at reduced costs, thereby making cryo-EM more accessible.

Many remaining challenges in structure determination by cryo-EM are associated with steps that precede cryo-EM imaging and involve protein purification, biochemical characterization and preparation of cryo-EM grids. Significant attention was drawn to the cryo-EM grid preparation where reproducible and controlled ice thickness is required [4,19]. Reduced protein stability in the thin film of the buffer before vitrification may cause protein denaturation [20] and often proteins adsorb at the air-water interface or support film, leading to preferred particle orientation [21,22]. Here also progress is being made by developing new grid supports and grid types [19,23–29] or faster plunging devices [30,31].

The analytic potential of single particle cryo-EM up to now has received less attention. It is generally recognized that in commonly practiced grid preparation by depositing microliter volumes of purified protein on EM grids followed by blotting with filter paper [32] less than one thousandth of the protein solution is retained on the plunge-frozen grid, whereas the rest is discarded together with the blotting filter paper. New cryo-EM grid preparation approaches that minimize the amount of protein solution deposited on EM grid are being developed. They include jetting picoliter droplets on the EM grid [30], pin printing [33,34], nanofluidic chips (25), or on-grid writing with a capillary (cryoWriter method) [35,36]. The cryoWriter approach was also used to isolate a target protein from cell lysate inside a capillary using superparamagnetic beads trapped by magnets [37]. This is a single example of when miniaturization was applied to the whole process of cryo-EM grid preparation, including protein purification. However, cryoWriter relies on a simple purification procedure, which allows only a single-step purification. It does not permit control of protein concentration often required for reproducible preparation of cryo-EM grids, and it requires an established purification procedure. This leaves a gap between this simple purification approach and more sophisticated multistep purification procedures requiring precise protein control at each step of the multistep purification process.

The common strategies of cryo-EM sample preparation start with hundreds of milliliters of cell extracts to end up with milligrams to micrograms of purified proteins, of which only a millionth to a billionth fraction is later imaged by cryo-EM. To reduce the gap between the amount of protein required to solve a high-resolution protein structure and the amount of starting biological material used for cryo-EM sample preparation, we developed a micro isolation (MISO) method that uses microfluidics to accomplish protein purification on chip in a continuous process and prepare cryo-EM sample from the purified protein using only tens of nanolitres of purified protein solution per EM grid. We demonstrate that the method can be used to solve cryo-EM structures starting from the cell mass equivalent to a single bacterial colony and for heterologously expressed mammalian membrane proteins from half a 10 cm dish, which is 100-500 fold reduction relative to the conventional purification schemes.

## Results

### Rational and implementation of MISO system

Assuming that ∼10^6^ particles need to be imaged and that the sample is prepared at a concentration of 0.01-10 mg/ml, the protein mass and volume of protein solution needed to solve the protein structure is in the order of picograms and picolitres. Consequently, volumes conventionally operated during the protein purification process can be scaled down by at least six orders of magnitude. Such miniaturization makes the manual liquid handling in between the individual steps impractical, which necessitates implementing all the steps involved in protein purification and cryo-EM grid preparation in a continuous process (**Figure 1a**). To realize this idea in practice, we developed a microfluidics chip and a device that facilitates on-chip protein purification and cryo-EM grid preparation (**Figure 1b-c, Supplementary** Figures 1**, 2**).

**Figure 1.**
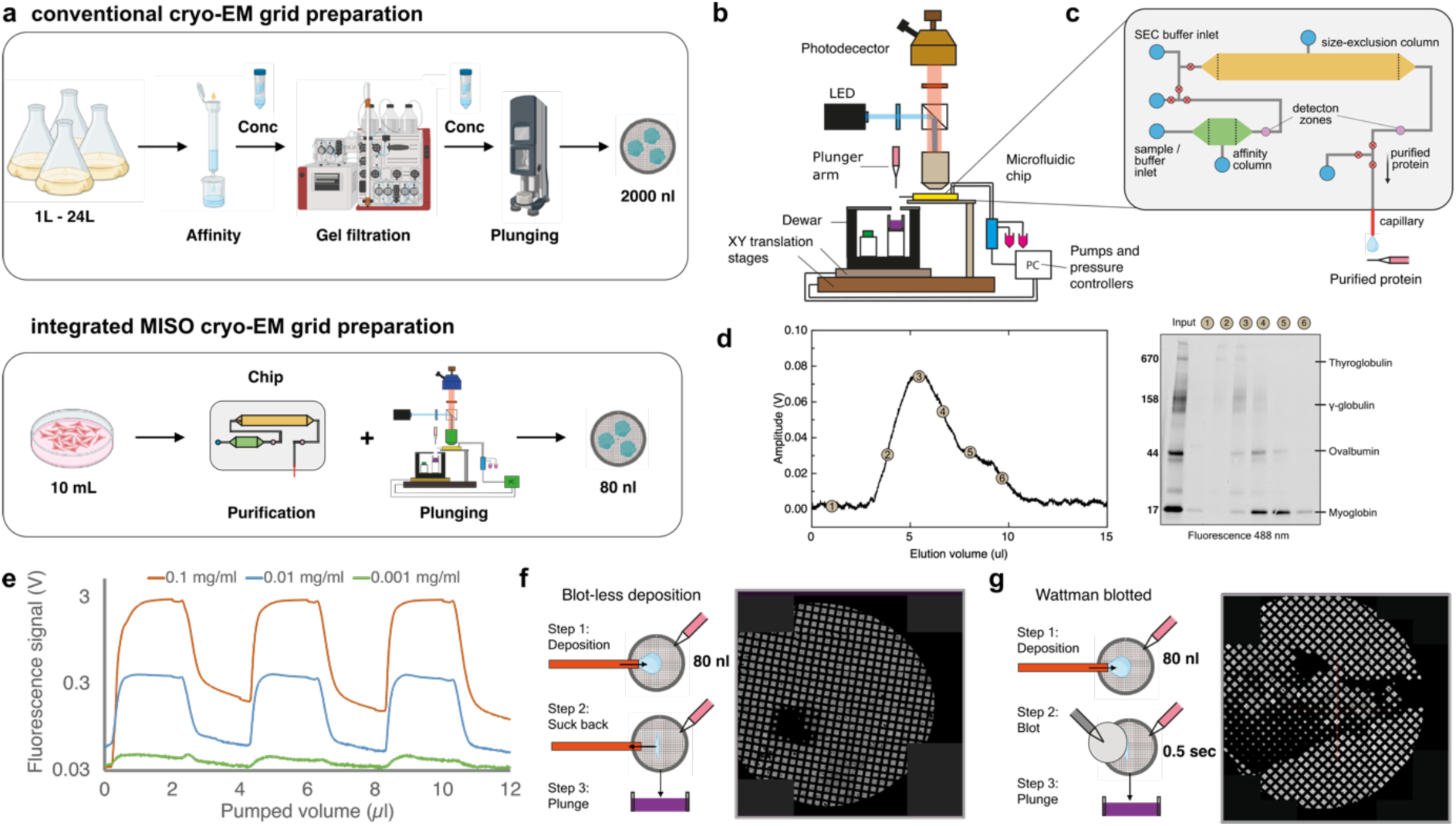
| **MISO principle and method characterization. a,** Conventional versus MISO workflows from cell extract to cryo-EM grids. **b**, Schematic of the MISO device. The key modules and elements are labelled. **c**, Schematics of a 2-column MISO chip. Red circles show position of on-chip pneumatic valves. **d,** Protein detection by fluorescence. Green fluorescent protein (GFP) can be detected to concentration of 1 µg/ml. **e**, Separation of Alexa488-labeled gel filtration standards on a 10 µl Superdex 200 Increase column. Proteins in the collected 1.5 µl fractions visualized using SDS-PAGE and fluorescent imaging (emission WL 488 nm). **f, g** Blot-less (f) and blotted (g) cryo-EM grid preparation approaches and resulting grid atlases.

We implemented an on-chip two-step purification process with affinity purification as the first and size-exclusion chromatography (SEC) as the second step. This covers purification protocols used for many water-soluble and integral membrane proteins. A microfluidic chip was designed and fabricated in silicone elastomer polydimethylsiloxane (PDMS, **Supplementary** Figure 2) using the standard process of photo-lithography and soft-lithography [38]. The chip contains a 0.5 µl column for affinity purification and a 5 µl column for size-exclusion chromatography (SEC). Due to photolithography constraints, the column’s height was ∼150 μm while its length and width were a few millimeters (**Supplementary** Figure 2). Filters were designed at the in- and outlet of the column to retain resin beads inside, and a filling port was used to fill columns with chromatography resins. Detection zones, with a volume of ca. 100 nl, were designed downstream of each column to facilitate protein detection. On-chip, slightly open doormat valves [39] enabled accurate flow control preventing cross-contamination of buffers and minimizing dispersion due to the parabolic flow velocity profile in the microchannels [40]. The purified protein was taken out of the chip through a fused silica capillary mounted horizontally directly into the microfluidic channel (**Figure 1c, Supplementary** Figure 2). The capillary reduces the dead volume to below 1 μl and consequently dispersion and dilution between the observation zone and a grid.

The chip was operated on a setup (**Figure 1b** and **Supplementary** Figure 1) that comprised a microscope for chip observation and fluorescence detection, a syringe pump and pressure regulators that controlled liquid flow in the chip, and operation of on-chip pneumatic valves. A plunger module comprised automated tweezers to hold and transfer the grid between the capillary and a cryogenic dewar. An automated arm with mounted filter paper was used for grid blotting during cryo-EM grid preparation (**Supplementary** Figure 1). The microfluidic chip, vials with protein extracts and the syringe pump were thermostatted to 4°C.

Due to the high absorbance of PDMS at 280 nm, a wavelength typically used for protein detection, the proteins on the chip were detected using fluorescence. We characterized the sensitivity of the fluorescence detection using green fluorescent protein (GFPuv, **Figure 1e**), for which the concentrations down to 0.001 mg/ml (37 nM of GFP, MW 27 kDa) were reliably detected. Such sensitivity is sufficient to detect proteins with a molecular weight of several kilodaltons at concentrations commonly used for cryo-EM (0.01-10 mg/ml) [4].

Unlike bulky chromatography columns, our on-chip columns were relatively shallow and wide (the dimensions of a 5 μl column were 20x2.5x0.15 mm); they were packed and ran at low pressure (below 0.2 MPa). To assess their capacity for separating macromolecules, we estimated the resolution of a ∼10 μl SEC column using common protein MW standards fluorescently labeled with Alexa Fluor 448 dye (**Figure 1f)**. The chromatogram of a mixture of these standard proteins resolved two peaks, and SDS-PAGE showed the separation of the proteins in the different elution fractions. We estimated the column plate number as 2000-5000 plates per meter. This is around 5 to 20-fold lower than the resolution of analytical HPLC columns (40,000 plates per meter) but sufficient for buffer exchange and separating large protein complexes from smaller proteins.

The volume of the elution peak from the MISO chip was in the order of a microliter. Direct coupling of the column outlet to the cryo-plunger permitted us to fractionate the peak by utilizing 20–80 nl of protein solution per EM grid. Two methods were used to prepare cryo-EM grids: (1) blotless deposition in which around 20 nl of protein solution was dispensed on the EM grid and sucked back creating thin areas at the periphery of the deposited drop (**Figure 1f**, **Supplementary Video 1**) or (2) by applying ∼80 nl volume on the grid, spreading it over the grid surface by moving the grid relative to the capillary followed by a brief grid blotting with a filter paper (**Figure 1g**, **Supplementary Video 2**). These two approaches were complementary to each other. The blotless application avoided contact with the paper and was successfully used to vitrify test protein samples (**Supplementary** Figure 3). However, it generated a limited number of squares suitable for data collection. The second approach, blotting cryo-EM grids with filter paper, consistently produced larger grid areas suitable for high-resolution cryo-EM data collection (**Supplementary** Figure 4).

### Validation of the micro purification and EM grid preparation workflow

We used *E. coli* cells overexpressing an established cryo-EM protein sample β-galactosidase (βG) [41] with engineered 6xHis-tag [42] to test the workflow of the developed MISO chip and the plunger. *E. coli* cytoplasmic extract was fluorescently labeled with Chromeo P503 dye that reacts with primary amines such that the formation of the adduct strongly enhances its fluorescence [43]. Consequently, only labeled proteins are fluorescent but not the unreacted dye. We verified that individual purified proteins and whole cytoplasmic extracts can be labeled and visualized using fluorescence imaging (**Supplementary** Figure 5).

Various affinity column volumes were tested by gradually decreasing the column volumes starting from 50 μl. Even though the photolithography allows manufacturing columns with volumes down to 100 pl, considering practical constraints such as protein adsorption on the channel surfaces (see below), the accuracy of dispensing, the possibility of analyzing eluted protein using SDS-PAGE, and negative stain EM, the smallest column volume implemented on the chips was limited to 0.5 μl. A second downstream 5 μl SEC column was added primarily for buffer exchange and separating the target protein from aggregates (**Supplementary** Figure 2a**, b**). The volume ratio of 1:10 between the affinity and size exclusion column allowed loading of the complete elution peak from the affinity column into the SEC for an efficient purification process.

βG purification was accomplished on a chip filled with Ni-NTA (0.5 μl) and Superose 6 (5 μl) resins. Initial experimental verification of the designed system was performed on *E. coli* cytoplasmic extract containing around 20 μg of βG (**Figure 2a**). The complete purification process included column and chip equilibration (∼100 μl), sample loading (20 μl), washing (120 μl), and elution (∼10 μl, **Figure 2b**). Affinity column washing with ∼60 μl was needed to stabilize the fluorescence baseline signal. This accounts for dead volume in the connecting tubing, dispersion and possibly adsorption of proteins on the microfluidic channel walls. βG was eluted from the 0.5 μl Ni-NTA column in a volume of around 1μl (**Figure 2c**), half of which was directed into the size exclusions column and eluted in a peak with a half-width of 1.0 μl (**Figure 2d**). It is noteworthy that the intensity of the size-exclusion peak was around 20 times lower than that of the affinity column elution peak. This cannot be explained by mere dispersion and is most likely caused by the unspecific protein adsorption on the channel and column walls and resin beads. The eluted protein solution was pure (**Figure 2e**) and was directed towards the capillary outlet, applied on cryo-EM grids, and plunge-frozen (**Figure 2f, Supplementary** Figure 4a).

**Figure 2.**
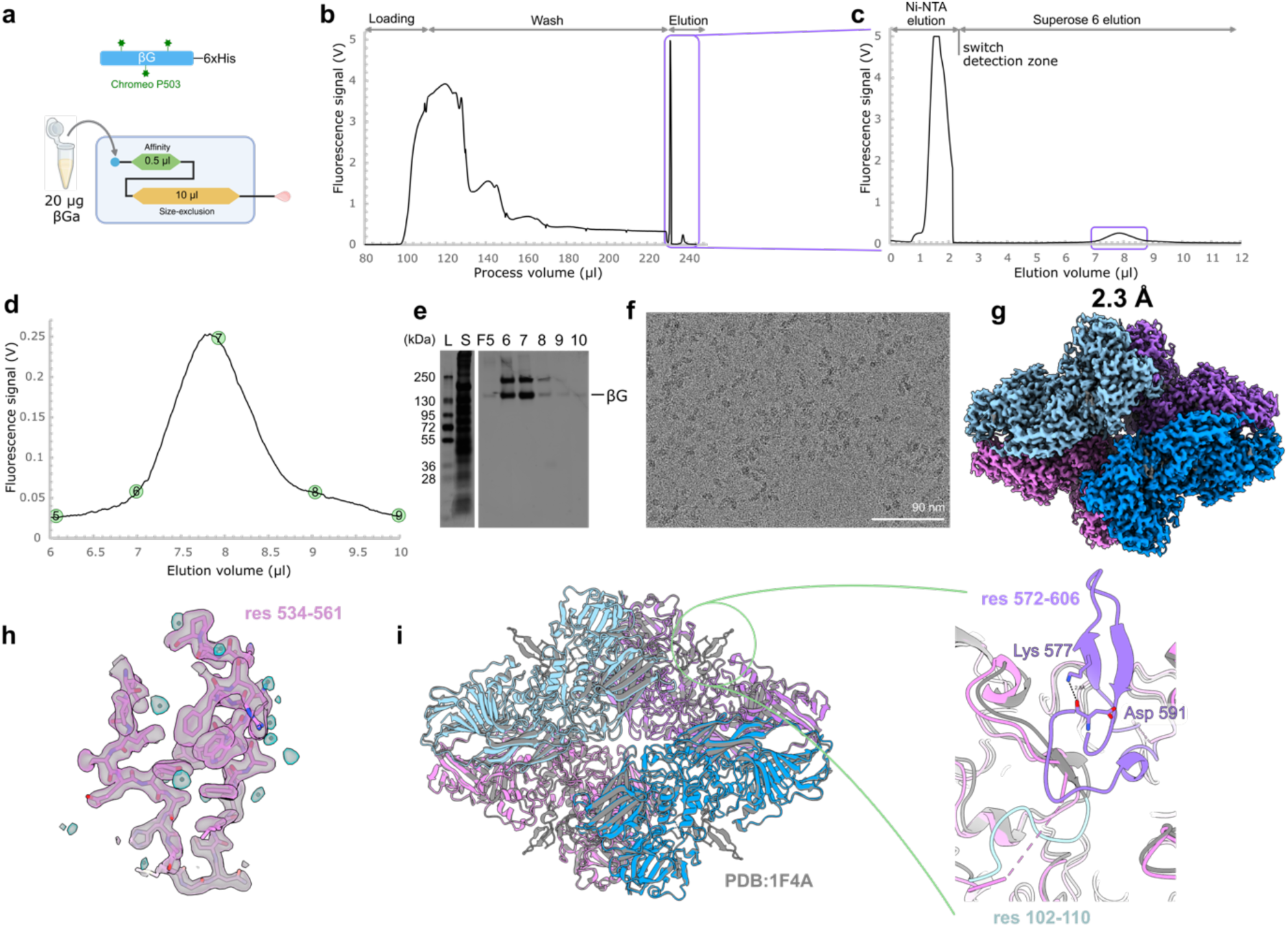
| Purification and cryo-EM structure of β-galactosidase isolated from 20 μg-containing cell extract. **a**, Experiment schematics. A 2-column MISO chip was used: 0.5 μl column filled with Ni-NTA resin and 5 μl column filled with Superose 6. **b**, Complete chromatogram of purification process shown starting from sample loading step. The initial system wash (first 80 μl) is omitted. Total cell extract was labelled with Chromeo P503 dye. **c**, Protein elution from Ni-NTA and SEC columns. **d**, Close up of SEC elution peak. Positions of the collected fractions are indicated by the numbered circles. **e**, Image of the SDS-PAGE with cytoplasmic extract (S) and fractions (F) collected from SEC column. **f**, An example of a micrograph from cryo-EM dataset collected using a holey EM grid and used for the map reconstruction. **g**, 3D cryo-EM map of βG reconstructed to 2.2 Å from a grid prepared using MISO device. **h**, Map details showing resolved side chains and ordered water molecules. **i**, Structural alignment between βG model built into map of Chromeo P503-labeled protein prepared using MISO and unlabelled crystal structure of βG (PDB: 1F4A). Zoom in on the disordered loops shown in purple and cyan. Lys 577 hydrogen bounded to backbone of Asp 591 are shown as sticks.

We initially used a blotless cryo-EM grid preparation, which resulted in usable cryo-EM samples (**Supplementary** Figure 3), but the reproducibility and the number of squares with optimal ice thickness were limited. Therefore, we switched to using at first manual and later automated paper blotting with 0.5 s blotting time. It reliably produced cryo-EM grids with tens of usable squares (**Supplementary** Figure 4). The elution peak was fractionated by plunging eight grids from the beginning to the end of the peak to scan a range of protein concentrations and particle sizes. Around 4,000 usable micrographs were collected from a single cryo-EM grid with the highest particle density and a cryo-EM reconstruction of βG was calculated to a resolution of 2.2 Å (**Figure 2g-h**, **Supplementary** Figure 6**, Supplementary Table 1**). The resulting 3D model was nearly identical to unlabelled βG solved by crystallography [44] (RMSD 0.41 Å for 3701 Cα atoms, **Figure 2i**). Loops, including residues 102-110 and 572-606 that form characteristic β-hairpin protrusions on the βG surface, were missing in the reconstruction obtained from the MISO chip (**Figure 2i**). The disorder is likely caused by the modification of Lys 577 by Chromeo P503 dye. In the crystal structure, Lys 577 stabilizes the loop conformation through a hydrogen bond to the backbone carbonyl oxygen of Asp 591 (**Figure 2i**). Examination of the other side chains for modifications did not reveal the fluorescent dye-compatible modifications even though additional densities were observed next to several His and Cys residues indicating their chemical modifications. In the cryo-EM reconstructions obtained with βG prepared by conventional methods and labeled with Chromeo P503 or Alexa Fluor dyes, the same loops were disordered supporting the chemical modification as the cause of the disorder.

### Structure of β-galactosidase from a single colony

βG has high expression levels when overexpressed in *E. coli*. The corresponding bands on SDS-PAGE can be observed in the crude cytoplasmic extract (**Figure 3e**). We were curious to see whether the traditional workflow of structure determination requiring large starting volumes of bacterial or eukaryotic cell cultures could be shortcut to solving the structure of βG from a single bacterial colony. To test this, an *E. coli* colony was scraped from a Petri dish, diluted in 1 ml of medium (**Figure 3a**), and induced with IPTG for four hours. The cells were lysed, and the cytoplasmic extract, estimated to contain around 1 μg of βG, was labeled with Chromeo P503 dye and purified on a MISO chip, as described in the previous section. The protein elution from the Ni-NTA column was reliably detected, however, our attempts to detect the elution peak after the SEC column were initially unsuccessful. We realized that protein losses were due to the protein adsorption on the hydrophobic microfluidic chip surface and SEC beads. To overcome this, we added 0.2% of detergent β-dodecyl maltoside (DDM), which improved the protein recovery after the SEC (**Figure 3c**). When comparing the chromatograms, the affinity elution peak intensity from the 20 μg experiment was more than 20 times lower compared to the 1 µg experiment, but from the SEC column, it was only around 3 times lower. Hence, using DDM improved βG recovery between the chromatography columns by around 6-fold. Cryo-EM grids with an additional thin carbon film were used to obtain satisfactory particle density on the EM grids to which βG purified from a single colony was applied (**Figure 3f**). A dataset comprising 5,700 images was collected from a single grid and a 3D map of βG was reconstructed to a resolution of 2.2 Å (**Figure 3g, h, Supplementary** Figure 7**, Supplementary Table 1**). The refined structure was identical to the one solved from 20 μg of starting βG amount.

**Figure 3.**
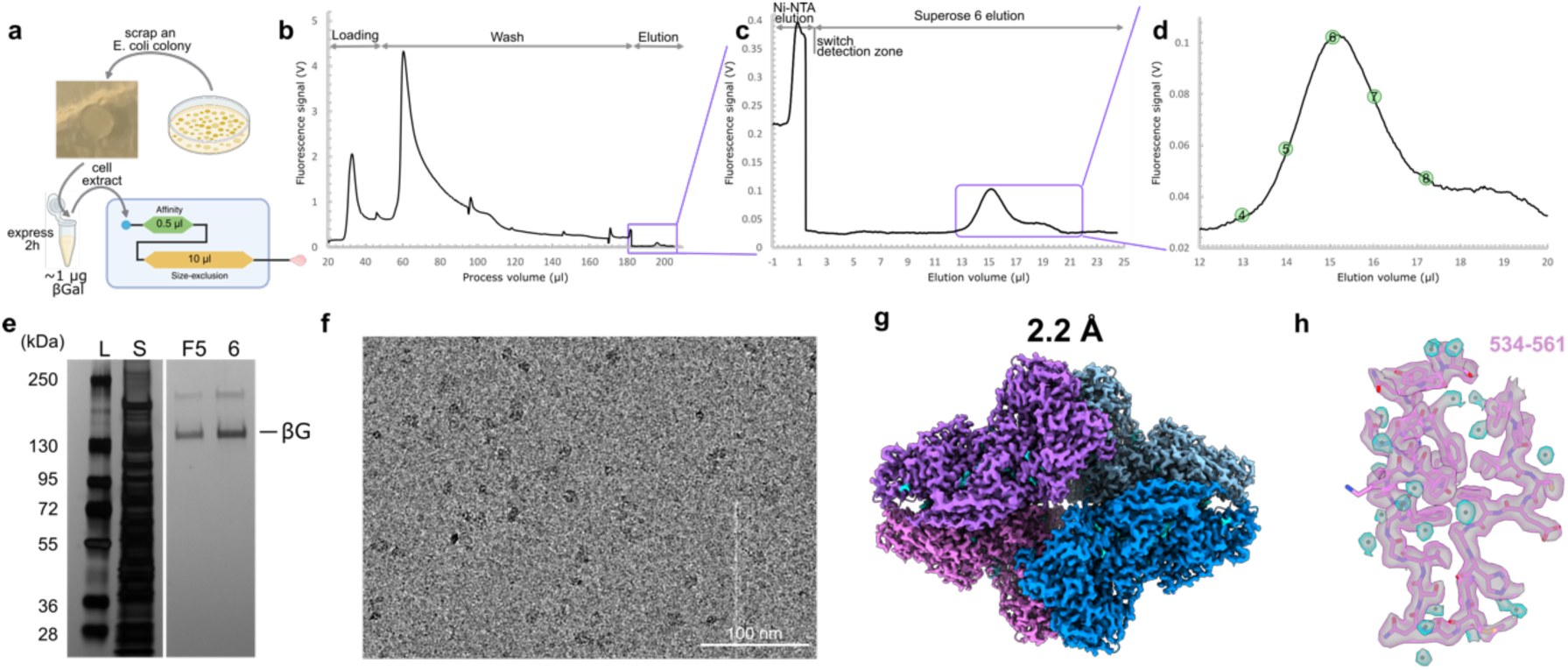
| **Structure of β-galactosidase solved starting from a single *E. coli* colony. a**, Schematics of the experiment. Purification chip was the same as in Figure 2. **b**, Chromatogram of the βG purification. Signal intensity at WL 593 nm from proteins labelled with Chromeo P503 dye. **c**, Close up view of the protein elution. **d**, Elution peak from 5 μl SEC column was fractionated into 1 μl fractions. **e**, SDS-PAGE from the purified βG. **f**, An example of micrograph obtained on graphene-oxide coated grids from βG purified starting from 1 μg βG in cytoplasmic extract. **g**, **h**, Cryo-EM map and map fragment of the reconstruction of βG purified from 1 μg containing cell extract.

This experiment shows that the 2-step purification procedures can be performed on the MISO chip and the eluent is sufficiently concentrated for preparing cryo-EM grids suitable for high-resolution single-particle structure determination starting from 1 μg of protein in a crude cytoplasmic cell extract, which in case of βG can be extracted from a single colony after protein expression. This demonstrates the advantages of MISO approach for accelerating cryo-EM structural studies.

### Membrane protein example 1: TMEM206, a proton-activated chloride channel

Next, we applied the MISO approach to three eukaryotic membrane proteins to validate it. These proteins are expensive and labor-intensive to obtain in pure form, yet they are highly relevant to our understanding of human cellular physiology and many are drug targets. Reducing the required amounts of expression media and cell mass is critical in facilitating structural work. As a first example, we used the bovine proton-activated Cl^-^ channel TMEM206/PACC1 (*bt*TMEM206) stably expressed in FlpIn T-Rex 293 cells with a C-terminally fused venus-YFP (vYFP) followed by a streptavidin binding peptide (SBP) tag. TMEM206 are homo-trimeric, glycosylated trans-membrane proteins with a molecular weight of ∼40 kDa per subunit. Several homologs of TMEM206 have been structurally and functionally characterized [45–47]. Although the natural function is still under debate [48], TMEM206 channels are considered promising drug targets to prevent cell death after ischemic stroke or in lysosomal physiology. The expression level of *bt*TMEM206 in the established cell line was estimated to be 4 µg per 10 cm petri dish. Before using the MISO method for purification and cryo-EM grid preparation, *bt*TMEM206 protein without vYFP, was expressed in 150 10 cm dishes and purified with streptavidin beads in batch followed by SEC on a 5/150 Superdex 200 increase column on an HPLC system (**Figure 4d, h**). The peak fractions from the SEC column were pooled, concentrated and plunge-frozen on holey carbon grids and the structure of the channel was determined to a resolution of 2.9 Å (**Figure 4i-j, l-n, Supplementary** Figure 8**, Supplementary Table 1**).

**Figure 4.**
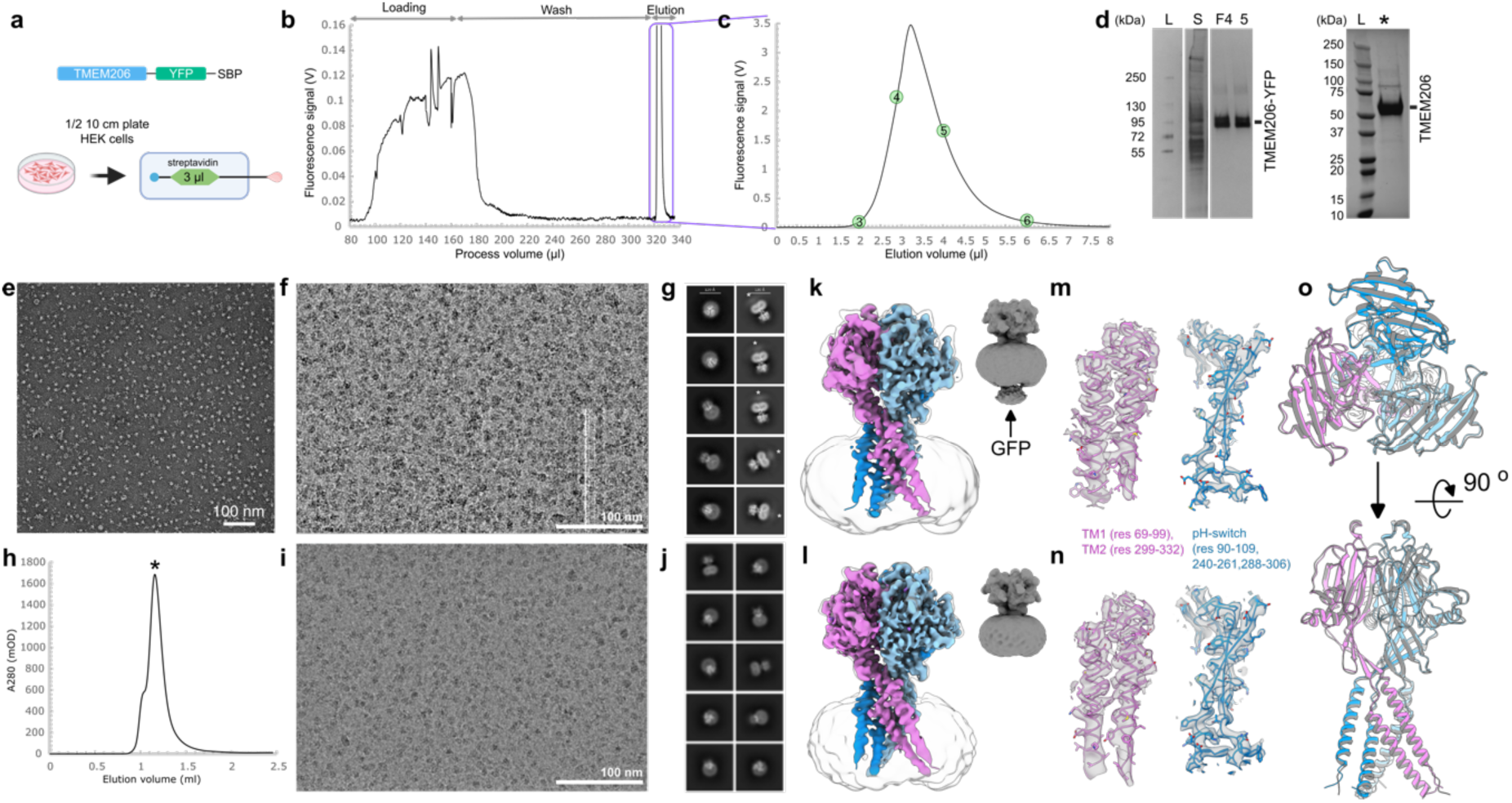
| **Preparation and structure determination of *bt*TMEM206 using MISO and conventional purification approach**. **a**, Schematics of the protein construct and MISO purification on a single 3 μl microfluidics streptavidin column. **b**, Complete MISO purification chromatogram of btTMEM206-YFP from cleared detergent-solubilized HEK cells. Fluorescent signal was detected at WL 525 nm and originates from YFP. **c**, Elution peak was fractionated into 1 μl fractions used for protein characterization. **d**, SDS-PAGE from *bt*TMEM206-YFP purified on MISO (left) and *bt*TMEM206 purified by conventional approach (right). The conventionally purified TMEM206 was gel-filtered (panel h) and the peak fraction was used for SDS-PAGE. **e**, Negative stain micrograph from MISO- purified sample shows homogeneous protein particles. **f** and **i**, Cryo-EM micrographs of MISO-purified *bt*TMEM206-YFP and conventionally purified TMEM206 on holey EM grids. **h**, SEC from *bt*TMEM206 purified by the conventional method. **g, j-n**, Comparison of 2D class averages and 3D reconstructions from *bt*TMEM206-YFP (g, k, m) and *bt*TMEM206 (j, l, n). Additional diffuse density from YFP in 2D classes is marked with white asterisk (panel g). **o**, Structural alignment between *bt*TMEM206 (grey) and *bt*TMEM206-YFP (coloured) models shows that YFP does not modify the structure.

To prepare cryo-EM grids using MISO method, we used a single 3 μl column packed with the same streptavidin resin as in the conventional approach. The protein was detected using vYFP fluorescence (**Figure 4a**). BtTMEM206-YFP eluted as a narrow (half-width 1.3 μl) single intense peak (**Figure 4b, c**), was pure (**Figure 4d)** and monodisperse as judged from negative stain EM micrographs (**Figure 4e**), in agreement with the previous size SEC run (**Figure 4h**). Therefore, we prepared cryo-EM grids using protein eluted directly from the affinity column. A cleared cell lysate from half of a fully confluent 10 cm cell dish containing an estimated 2 μg of *bt*TMEM206 was purified on the 3 μl column. The eluted protein was directly plunge-frozen on holey cryo-EM grids (**Figure 4f, Supplementary** Figure 9). Similar to the βG procedure, the elution peak was fractionated by plunging eight grids. At the peak of elution, the protein concentration was sufficient for producing gold UltraAuFoil holey grids with high particle density comparable to the conventional protein preparation (**Figure 4f, i**). A dataset containing 7,300 usable images was collected from a single grid (**Supplementary** Figure 9) and a 3D reconstruction was calculated to a resolution of 3.0 Å (**Figure 4g, k, m, Supplementary** Figure 10**, Supplementary Table 1**).

The cryo-EM grids, 2D classes, and 3D reconstructions were very similar between *bt*TMEM206 solved using a conventional approach and TMEM206-YFP using MISO (**Figure 4**). A diffuse density on the cytoplasmic side consistent with vYFP connected by a flexible linker to the channel was observed in the 2D classes of the MISO samples (**Figure 4g**) and as a low-resolution density in the 3D reconstruction (grey insets in **Figures 4k** and **l**). The differences between *bt*TMEM206 and *bt*TMEM206-YFP structures were minimal (RMSD 0.75 Å for 763 Cα atoms, **Figure 4o**), indicating that fused vYFP does not change the TMEM206 structure.

This example demonstrates that the MISO method can be successfully applied to solving the structure of a 120 kDa membrane protein starting from 300-fold smaller amounts of eukaryotic membranes.

### Membrane protein example 2: TMEM16F, a Ca^2+^-activated lipid scramblase

Next, using the MISO approach we set out to determine a structure of a membrane protein that has smaller extra-membranous domains compared to *bt*TMEM206, lower symmetry, and for which we did not have an established cryo-EM grid preparation procedure by conventional means in our laboratory. We used a murine orthologue of a Ca^2+^-activated lipid scramblase belonging to the TMEM16 family (*m*TMEM16F). The TMEM16 family constitutes Ca^2+^-activated lipid scramblases and chloride channels that play important roles in the regulation of lipid asymmetry and initiation of subsequent signaling events, as well as the contraction of smooth muscles and chloride secretion. TMEM16A is considered a major drug target for the modulation of chloride secretion in lung epithelia in cystic fibrosis patients and is upregulated in different types of cancer. TMEM16F is required for the Ca^2+^-dependent exposure of phosphatidylserine to the outer leaflet of the plasma membrane [49], an essential step in the blood coagulation cascade in platelets, among numerous other processes linked to the breakdown of lipid asymmetry [50]. Recently, TMEM16F was linked to the pathology of COVID-19 and its druggability has been demonstrated [51].

Murine TMEM16F is a membrane-embedded dimer with a dimeric MW of 210 kDa and comparably small cytoplasmic domains [52,53]. A construct of *m*TMEM16F with C-terminal vYFP and SBP-tag was stably and inducible expressed in FlpIn Trex 293 cells. A cell pellet from a single 10 cm dish was used for structure determination (**Figure 5**). The purification and cryo-EM grid preparation was performed using the protocol described in the previous section. The protein was purified on a MISO chip with a 3 μl streptavidin agarose resin (**Figure 5a**), and the structure of mTMEM16F-YFP was solved with only two rounds of experiments. For each experiment, we used the cell pellet of a confluent 10 cm dish of the induced cells that correspond to an estimated number of 10 mio cells and an amount of expressed *m*TMEM16F of 1-2 μg (as compared to the yields of other membrane proteins routinely purified in our lab). First, *m*TMEM16F was purified and eluted as a sharp 0.9 μl peak (**Figure 5b, c**). The eluted protein was highly pure and monodisperse (**Figure 5d, e**). Next, the purification was repeated, and the eluted protein was plunge-frozen on holey carbon and UltraAuFoil cryo-EM grids. The screening identified a grid with sufficiently high particle density and ice thickness suitable for cryo-EM data collection (**Figure 5f, Supplementary** Figure 4). 12,500 micrographs were collected from the grid and a map of *m*TMEM16 was reconstructed to a resolution of 3.5 Å (**Figure 5 g-j**, **Supplementary** Figure 11, **Supplementary Table 1**).

**Figure 5.**
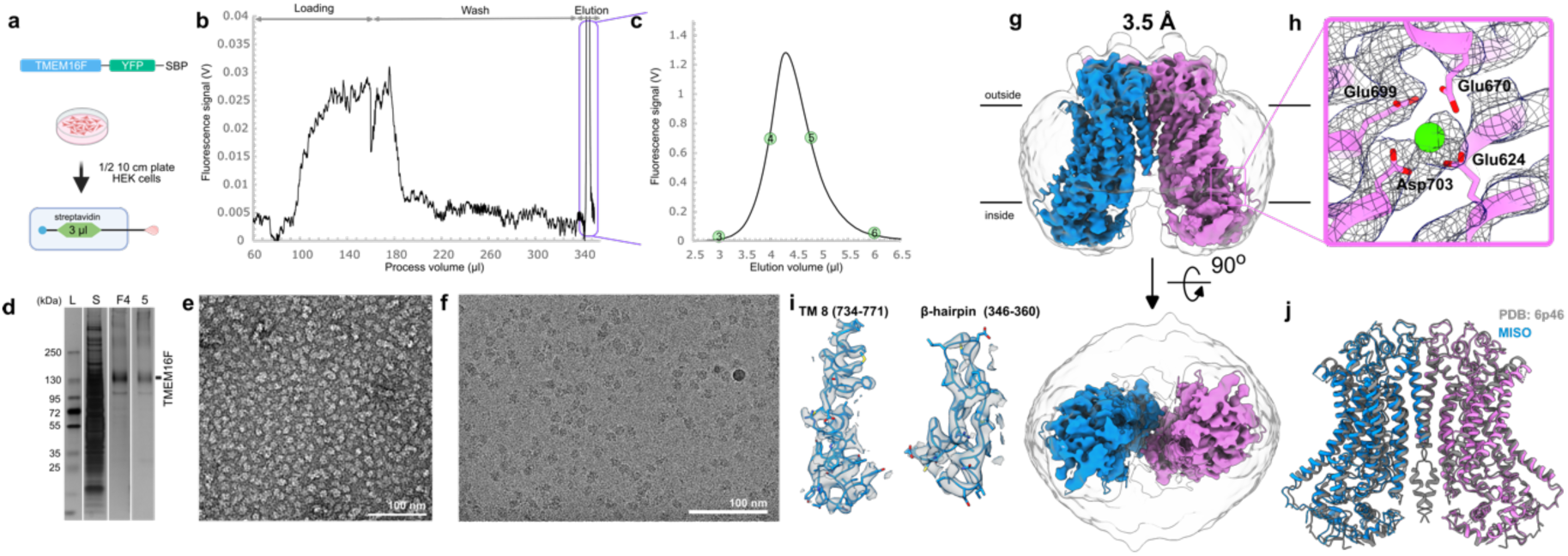
| De**termination of *m*TMEM16F structure using half a 10 cm dish of HEK cells. a**, Schematics of the protein construct and MISO purification on a single 3 μl streptavidin column. **b**, Complete MISO purification chromatogram from a cleared lysate of HEK cells. Fluorescent signal from YFP was detected at WL 525 nm. **c**, Elution peak was fractionated into 1 μl fractions used for protein characterization. **d**, SDS-PAGE from *m*TMEM16F-YFP purified on MISO. **e**, Negative stain micrograph from MISO-purified sample shows homogeneous protein particles. **f,** Cryo-EM micrographs of MISO-purified *m*TMEM16F-YFP on holey EM grids. **g,** 3D reconstruction of TMEM16F-YFP to a resolution of 3.5 Å . High-resolution map is overlayed with a low-pass filtered map to show the detergent micelle. **h**, Close-up view on the Ca^2+^ binding site with bound Ca^2+^ ion. **i**, Details of density map for a trans-membrane helix and for a β-hairpin. **j**, Structural alignment between MISO purified *m*TMEM16F-YFP and *m*TMEM16F prepared by conventional approach (PDB 6P46).

The model built into the density was very similar to the reported structure of mouse TMEM16F (**Figure 5j**, RMSD 1.25 Å over 1215 Cα). Even though calcium was not added to the buffer, the ambient concentration of calcium present in the buffers containing no chelator agents (typically micromolar amounts) was sufficient to bind one Ca^2+^ ion per binding site (**Figure 5** **h**).

This example demonstrates that new structural projects can be conducted entirely relying on the downscaled purification and cryo-EM grid preparation protocols. A single 10 cm dish of HEK cells and only 2 MISO chips were sufficient to obtain a structure of the small eukaryotic membrane protein.

### Example 3: Protein identification from a cryo-EM map

The examples described up to now dealt with the determination of the structures for proteins with known identity. In this example, we describe another case of a membrane protein for which structure determination using MISO allowed us to identify the protein from a reconstructed cryo-EM map. We used a pellet of insect Sf9 cells in which a membrane protein of interest was expressed as a fusion with eGFP and a twin-Strep tag (**Figure 6a**). We asked not to reveal the protein identity but did use an established protocol of protein purification as a starting point. First, the equivalent of 50 ml of Sf9 cell culture was solubilized in LMNG and, after ultracentrifugation, loaded on a 10 μl streptavidin MISO column, washed and eluted as a ∼3 μl wide peak fractionated into 3 μl fractions (**Supplementary** Figure 12a). A silver-stained SDS-PAGE displayed an enriched band with a molecular weight of around 150 kDa (**Supplementary** Figure 12a) that co-eluted with many other less intense, presumably contaminating proteins. To improve the purity, 60 μl of solubilizate was filtered through a 0.22 μm filter and purified on a 1 μl MISO column using more extensive washing (**Supplementary** Figure 12b). The protein eluted as a sharp ∼1 μl peak which displayed higher purity and contained monodisperse particles of ∼160 nm in diameter when visualized by negative stain EM (**Supplementary** Figure 12e, micrograph a).

**Figure 6.**
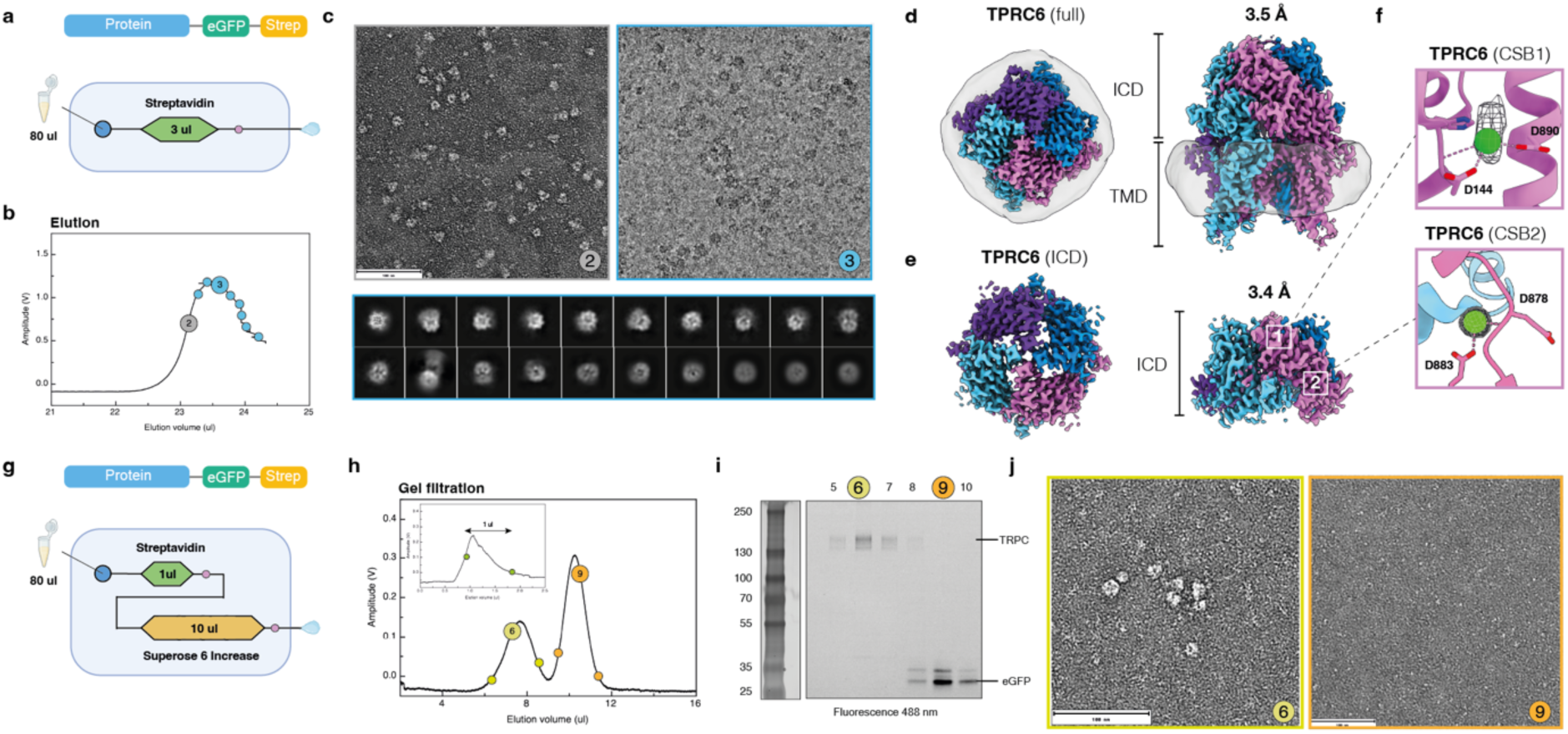
|Purification and identification of TRPC6 channel. **a**, Protein constructs and schematics of the experiment. 80 μl of extract was purified on a 3 μl streptavidin column. **b,** Elution peak detected at WL of 525 nm was fractionated into 1 μl fraction used for negative stain EM (grey in the chromatogram) while subsequent 80 nl fractions were spotted on holey gold grids (blue) and cryo-plunged. **c**, Negative stain micrograph and an example of cryo-EM micrograph from MISO device. Panel below shows 2D classes from the cryo-EM dataset. **d,** 3D cryo-EM maps of the of the complete membrane protein complex solved to resolution of 3.5 Å. **e,** 3D reconstruction of intracellular domain (ICD) only resolved to 3.4 Å. **f,** Density for two distinct Ca^2+^ ions bound to calcium binding sites (CBS) with fitted structure of human TRPC6 (PDB 6UZ8). **g,** Experimental scheme for purification of TRPC6 on a 2-column MISO chip. **h,** Protein purified on 1 μl streptavidin column was further polished on a 10 μl Superose 6. **i,** SDS-PAGE imaged using fluorescence at WL 488 nm. **j,** Negative stain EM micrographs from the selected fractions.

To prepare cryo-EM grids, a 3 μl streptavidin column was loaded with 80 μl of solubilizate, corresponding to ∼ 2.5 μg of the protein of interest (**Figure 6a**). The column volume was increased to minimize dilution of the elution peak between the column and the grid where the dead volume was ∼0.7 μl. Eight grids were plunged from ∼1 μl elution peak (**Figure 6b**) and two fractions were collected for negative stain EM (**Figure 6c**). Screening of the cryo-EM grids identified a grid suitable for data collection (**Figure 6c, Supplementary** Figure 13) from which 6,572 movies were collected. After processing and 3D classification, two maps were reconstructed (**Supplementary** Figure 14**, Supplementary Table 1**). One map reconstructed to 3.5 Å resolution resolved the transmembrane region with a detergent belt and a large solvent-exposed domain. In contrast, the second map obtained at a resolution of 3.4 Å corresponded to the solvent-exposed domain only (**Figure 6d, e**). A *de novo* model was built using ModelAngelo [13], which generated 151 polypeptide fragments with lengths of up to 166 amino acids. Hidden Markov model sequence profiles of the fragments were used for HMMER search [54] that consistently identified short tetrameric transient receptor channel C6 (TRPC6) as the top hit (**Supplementary** Figure 15). Due to the high sequence identity of TRPC6 between the organisms, the exact organism identity remained ambiguous. We fitted and refined the structure of human TRPC6 into the reconstructions (**Supplementary** Figure 15). The resulting model had densities consistent with bound metal atoms in two Ca^2+^-binding sites (**Figure 6f**), and the overall model conformation was consistent with TRPC6 in a Ca^2+^-bound conformation with RMSD of 0.79 Å (**Supplementary** Figure 15a).

We further analyzed the TRPC6 channel using a 2-column MISO chip to see if the quality of the cryo-EM sample can be further improved. Using a combination of 1 μl strep and 10 μl Superpose 6 Increase columns, we were able to separate two fluorescent peaks on the SEC column (**Figure 6g, h)** which after separation on SDS-PAGE followed by fluorescence imaging and silver staining revealed two bands corresponding to the TRPC6 monomer (∼150 kDa) and what is likely to be a cleaved eGFP (∼30 kDa) (**Figure 6i, j**). This demonstrated that the 10 μl SEC column was able to improve the protein purity. No void volume peak typically observed during the large-scale TRPC channel purifications was resolved suggesting that rapid, on-chip purification might be gentle enough to minimize protein degradation and the formation of larger aggregates.

This example demonstrates the potential of MISO purification for performing protein isolation, characterization, and cryo-EM sample preparation for proteins with unknown identity and indicates an MISO potential for becoming a tool for the discovery and structural characterization of protein complexes.

## Discussion

The core advantage of the described MISO method is its capacity to operate with volumes and protein quantities that are three orders of magnitude smaller than those used in conventional protein purifications. Consequently, the whole cryo-EM sample preparation workflow, starting from cell extracts to cryo-EM grid plunging, requires hundreds to thousands of times less biological material. This corresponds to around 1 μg of starting amount of target protein in protein extracts that can be obtained from a single *E. coli* colony or less than 10 ml of HEK cell culture (corresponds to one 10 cm dish). The purification and plunging are completed within a few hours after cell harvesting while being compatible with commonly used multistep protein purification strategies.

Here, we demonstrated a proof of principle with a soluble bacterial protein and three eukaryotic membrane proteins. Because MISO uses a few hundred to thousand times fewer starting materials than conventional purification methods, culture scale-up is not needed before purification, which reduces the time associated with cryo-EM grid preparation. In addition, MISO miniaturization opens the possibilities for structural studies that were previously too challenging or too resource-intense, such as protein structure determination of native proteins or native protein complexes from primary cells and small amounts of animal tissues, including biopsies.

Protein elutes from the MISO chips in a volume of around one microliter. This volume is sufficient to assess protein purity by SDS-PAGE, integrity by negative stain EM, and to optimize the purification protocol. A combination of affinity chromatography with SEC, a commonly used purification strategy to prepare cryo-EM grade protein, was implemented on MISO chips. Importantly, however, we observed that a single affinity column was sufficient for obtaining cryo-EM grade protein. When eluted from a 3 μl streptavidin column, the tested membrane proteins were concentrated enough to yield adequate particle density on holey grids, and no aggregates, commonly separated in the void volume of SEC, were observed in the cryo-EM micrographs. This may stem from the rapidity of the purification and, consequently, better preservation of proteins or from omitting commonly used spin concentrators that create high local protein concentrations that may lead to aggregation.

The elution peaks from MISO chips were fractionated into eight fractions allowing to sample various protein concentrations and sample heterogeneity depending on the retention volume. Compared to blotted sample preparation, blotless grid preparation did not provide obvious advantages when tens of nanoliters of protein were deposited on an EM grid. Because the applied volumes were smaller than those used for conventional cryo-EM grid preparation, the blotting time was reduced to hundreds of milliseconds minimizing contact of protein with the blotting paper and reducing potential air-water interface effects [21].

The microfluidics approach required designing a continuous process where all the purification steps and cryo-EM grid preparation are part of a non-divisible workflow. Three design features were critical for the successful implementation of the MISO method: (1) minimization of dead volumes that reduce dispersion and non-specific protein adsorption, (2) implementation of on-chip valves enabling precise flow control and preventing buffer mixing, and (3) on-chip protein detection to ensure sufficient washing, optimal timing for protein injection into the SEC column and determining the position of elution peak for cryo-EM grid plunging.

Proteins on MISO chips were detected using fluorescence. We used either unspecific labeling by fluorescent dyes conjugated to surface-exposed primary amines or proteins that were fused to genetically encoded fluorescent proteins via flexible linkers. Non-selective protein labeling has the advantage of monitoring all proteins, including contaminants, during purification. However, we observed that modifications of lysins with dye, induced local loop disorder in βG (**Figure 2i**) and therefore should be used with caution. Genetically encoded fluorescent tags, on the other hand, did not alter protein structures and allow to quantification of the target protein.

Future improvements to the MISO method may focus on increasing the resolution of SEC microcolumns and developing an on-chip protein concentrating method that will extend the MISO approach to purifications on low-capacity affinity media. Systematically addressing protein adsorption in the microfluidic chip [55,56] will improve protein recovery and allow further scale-down of the MISO process. We observed that MISO purification required extended column washes. This can be due to a combination of dead volumes of connecting tubing that are an order of magnitude larger than the column volumes, and protein adsorption on the tubing and channel walls. Both will be investigated in the future.

The described MISO method scales down the purification process hundred- to thousand-fold and operates with sub-microgram protein amounts. Protein amounts applied per EM grid using MISO are in the order of 10 ng, which corresponds to ∼ 10^10^ molecules. Hence, a further potential downscaling of protein purification by three to four orders of magnitude to reach the theoretical limit of ∼10^6^ particles is possible. In this limit, protein must be concentrated to a volume of 0.1-100 pl (5-50 μm diameter drop) and spread over several grid squares. At such a small scale, the surface effects, including non-specific adsorption will present a major challenge [25], whereas protein dispersion and diffusion will become significant, and protein detection may be impeded by local heating and bleaching. Despite these difficulties, we envisage that scaling down the MISO approach by an additional one to two orders of magnitude is feasible. This will open up exciting possibilities to perform single particle cryo-EM on environmental samples, primary cell lines, organoids and small animal or human biopsy samples.

## Data availability

The protein models with the PDB accession codes 1F4A, 6P46, 6UZ8 were used in this study. The newly generated cryo-EM density maps and refined atomic models were deposited in the PDB and EMDB databases under accession codes: βG from 20 μg 9HPL, EMD-52333; βG from 1 μg 9HPM, EMD-52334; *bt*TMEM206 conventional 9HQN, EMD-52344; btTMEM206-YFP MISO 9HQO, EMD-52345; mTMEM16F-YFP 9HQP, EMD-52346; TRPC6 EMD-52486, EMD-52487

## Acknowledgements

We thank Dr. Piotr Kolata for assistance with assembly of MISO device, Dr. Raf Claessens for advising on the design of light detector, Dr. Marcus Fislage for assistance with cryo-EM data collection. We would like to acknowledge the funding provided by Vlaams Instituut Voor Biotechnologie, Fonds Wetenschappelijk Onderzoek (Grant Nos. G0H5916N, G054617N to R.G.E.), and by the European Research Council (Grant No. 726436 to R.G.E).

## Author contributions

G.E. developed the microfluidic chip, instrument, and software. G.E. and R.G.E. designed and constructed the plunger device. G.E. characterized and optimized the operation of MISO chips and plunger.

G.E. and S.D.G. characterized the MISO chip. A.S. fabricated MISO chips and prepared *E. coli* cells expressing βG. S.S. and J.D.B. designed the constructs for TMEM206 and TMEM16F, generated the stable cell lines, and established purification conditions for *bt*TMEM206 and *m*TMEM16F. B.S. and P.E. provided cells and purification protocols for TRPC6. S.S. and J.D.B. purified and plunged btTMEM206 using the conventional approach. G.E and S.D.G optimized MISO purification protocols. G.E. performed MISO experiments with β-galactosidase, *bt*TMEM206 and TMEM16F. S.D.G. performed MISO experiments with TRPC6. G.E., S.D.G. and J.D.B. collected and processed cryo-EM data. S.D.G. built, refined, and validated atomic models. R.G.E. prepared the original manuscript draft. R.G.E., G.E., S.D.G., S.S., and J.D.B. prepared figures, reviewed, and edited the manuscript. R.G.E conceived, managed, supervised the project, and acquired funding.

## Methods

### Experimental setup

The MISO setup consists of three modules: (1) the microscope with a microfluidic chip mounted on motorized stage, illumination system and photodetector for detection of the fluorescent signal, (2) microfluidic flow control system that includes a syringe pump, pressure controllers and valves, (3) cryo-plunger comprising a custom made dewar with thermostatted ethane vessel [42] and a motorized EM grid plunger arm, humidity chamber and blotting arm. The parts of the setup that come in contact with protein solutions are temperature controlled. Automation of the processes is achieved using a custom-built LabVIEW code. Schematics of the experimental set-up used is shown in **Figure 1b** and its implementation in **Supplementary** Figure 1.

### Microscope module

A Nikon Cerna microscope (CEA 1500, Thorlabs) fitted with 20X Mitutoyo Plan Apochromatic objective (MY20X-804, Thorlabs) and with 470 nm, 3 W power LED (SOLIS-470C, Thorlabs) in epi-fluorescence configuration with excitation filter at 469 nm (± 17.5 nm) was used to illuminate the detection zones. Depending on the fluorophore (Chromeo P503/GFP), the resulting fluorescence emission was monitored using a dichroic mirror with pass band greater than 569/505 nm and the emission filter at 593/525 nm (± 20 nm) onto a Silicon photodiode (SM1PD1A, Thorlabs). The signal was amplified using a custom-built biasing and two-stage amplification module based on AD8618 precision amplifiers and recorded using a National Instruments Data Acquisition board USB-6001. The data collected by the acquisition board were passed onto LabView2020 v20.0.1f1 and converted into digital chromatogram plots. The microfluidic chip was mounted on motorized stage (PLS-XY, Thorlabs) to align the detection zone to the focus of the objective lens in XY directions.

### Microfluidics flow control system

The flow in the microfluidic chip was controlled using a programmable syringe pump (P100-L, LSPone, Advanced Microfluidics SA) with 10 port distribution valve (V-D-1-10-050-C-U) and 25 µl syringe (S-25- P). This facilitated easy switching across various buffers and samples with minimal carry-over volume (2.8 µl). The valves on the microfluidic chip were regulated using Fluigent pressure controllers LineUp Link (LU-LNK-0002), Flow-EZ (LU-FEZ-2000), Push-Pull (ELUPPU1000), P-Switch (ELUPSW2000) and 2 ml high pressure P-Caps (P-CAP2-HP-PCK).

### Cryo-plunger module

The plunging system consists of four parts: plunging arm, cryo dewar, humidity chamber, and a plunger positioning system. The plunging arm is composed of a Dumont self-closing precision tweezer (N7, 0.03 mm), a low-profile solenoid (2EC), and a bipolar stepper motor (8HS13-0604S, StepperOnline). The tweezer was mounted on the axis of the stepper motor perpendicular to handle enabling circular motion of the mounted grid. The low-profile solenoid was used to open the tweezer for grid mounting and release (**Supplementary Video 2**). The rotation of the stepper motor, managed by a digital stepper driver (DM320T, StepperOnline) was controlled by Ardunio Uno.

The cryo dewar consists of a bath of liquid nitrogen in which a liquid ethane vessel and a slot for placing the grid box are immersed. The ethane chamber was made in copper, was thermo-isolated, and had a resistive heater mounted on the bottom and a thermocouple of type K to maintain constant temperature [42]. The temperature of liquid ethane was controlled by the Arduino Uno board that communicates with LabVIEW.

The plunger module was mounted on a platform rigidly attached to the microscope XY stage. The plunger arm and the dewar were placed on another XY stage controlled by a piezoelectric inertia motor (PD1/M, Thorlabs) which was used for precise positioning of the grid relative to the tip of the outlet capillary.

A humidity chamber was used during cryo-EM grid plunging. It was placed around the capillary and plunger arm and filled with humid air generated by ultrasonic humidifier. The blotting was done with the help of an arm with a mounted stripe of filter paper (**Supplementary** Figure 1). The arm position was controlled by a servo motor and Arduino Uno.

### Temperature control system

The temperature of the components of the experimental set-up that come in contact with protein solution including the microfluidic chip, syringe of the pump and the vials containing the protein and buffers was controlled by a water recirculation system in which water was typically cooled to 4°C by a Peltier thermoelectric cooler (AC 162, TE Technologies) and pumped by a Seal-less Coupling Centrifugal Water Pump (RS, Cat. Nr.: 817-0772).

### Microfluidic chip design

Two microfluidic chip designs were used: one-column and two-column chips (**Supplementary** Figure 2). The key elements of the MISO chip are columns designed as flat compartments with a height around 150 μm, a width of 2.5 mm and length of up to 20 mm (**Supplementary** Figure 2a**, c**). Pillars of 10 μm height that served to retain chromatography resin beads inside the column were introduced at the inlet and outlet of the column.

The two-column chip consisted of an affinity column (AC) with an inlet for sample/buffer introduction. The outlet of AC was connected to a detection zone-1 (DZ1, **Supplementary** Figure 2). The outlet of the DZ1 was connected to waste outlet-1 (WO1) and size-exclusion column (SEC). The WO1 was connected to waste collection vial via an external on/off valve. The on-chip-valves along with this external valve controls the direction of flow of fluid coming from the AC and could be directed either towards the waste vial or the size-exclusion column. The SEC has two inlets; one inlet directs the affinity purified protein to the SEC and the other directs equilibration buffer to SEC. The outlet of the SEC is connected to the DZ2. After DZ2, the microfluidic channel bifurcate. The eluate could be either diverted into the capillary for subsequent deposition onto the cryo-EM grid or to the waste outlet WO2. A separate column filling (CF) inlet was used for introducing the beads into the columns.

### Microfluidic chip fabrication

The MISO chips were made using PDMS (SYLGARD 184) through soft lithography and replica moulding [38]. The chips comprise three layers: the functional top layer, the thin PDMS membrane in the middle, and valve control in the bottom layer. The valve control layer was further supported by a glass slide.

The master mold for the functional layer was fabricated using 2-layers with different heights of columns and bread filter pillars. The design pattern was printed on a transparency mask with a resolution of 6 μm (Selba SA) and transferred on a silicon wafer coated with photoresist. The photoresist (SU83010, Kayaku Advanced Materials) was spin-coated at 3,000 rpm for 40 s using a Polos spin-coater (SPIN150i, SPS-Europe B.V.) to achieve a thickness of 10 um, followed by a soft-bake step on a hot plate at 95° C for 6 min. Then the pattern was transferred by illuminating the photoresist through the photomask with UV light at 365 nm and power of 30 W for 8 s using mask aligner UV- Kube 3 (Kloe). The post-bake step was performed by placing the wafer on the pre-heated hot plate at 95° C for 2 min. Then the unexposed areas were dissolved with SU8 developer solution (Kayaku Advanced Materials) for 10 min. During this step, the developer solution was continuously gently mixed on a shaker to ensure the removal of unexposed material. SU8 3050 was spin coated at 1500 rpm for 40 s on this master mold to achieve a thickness of about 100 μm. Soft baking was done at 95° C for 40 minutes. Another similar design pattern (without pillars in the mask) was used to expose the master mold at 30 W power for 9 s and post bake was done at 95 ° C for 5 min and developed it for 10 min. The resulting master was examined by an optical profilometer (Proflim3D, Filmetrics) to verify the structure details and the resist thickness. The thickness in the region of pillars was about 13 μm, whereas the columns and channels were at an average thickness of about 150 μm.

The master mold for the valve control layer was fabricated in a similar way as that of the second step of the above photo lithography process but with the use of an appropriate mask that contains the valve control design.

PDMS was mixed with curing agent in a ratio of 10:1 poured onto the master molds and heated in an oven at 60° C for at least 8 h. A thin PDMS membrane of about 25 μm thick was made by spin coating a mixture of PDMS and the curing agent in the ratio of 14:1 at 1000 rpm for 5 min onto glass slides that were silanized for 1 h. The coated PDMS membranes were heated at 60° C in oven for at least 8 h to complete cross-linking.

The valve control layer and functional PDMS layers peeled from its master mold were punched with biopsy punches 2.5 mm and bonded with the PDMS membrane after treating both the layers with air plasma treatment (100 W, 50 kHz, 1 min Femto Science, Cute). This combination was further bonded with the functional layer that had holes already punched in it using similar air plasma settings in due alignment under the microscope. This combination (with through holes for the inlets and outlets) was bonded with a clean glass slide using air plasma treatment.

The channels of the bonded chips were cleaned using isopropyl alcohol to remove the trapped air and subsequently, the column(s) were filled with beads through the column-filling inlets. The selected chromatography resin was filtered through an EASYstrainer filter (70–100 µm, Greiner Bio-One) to prevent channel blockages. A microfluidic flow controller (Flow EZ 1000 mbar, Fluigent) was used to flow the bead suspension in the columns via the inlets. To achieve uniform packing of the resin, the pressure was gradually increased from 150 mbar to 1000 mbar. The flow was stopped once the resin reached the inlet, and after pressure equilibration, the tubing was cut near the inlet and sealed with a two-component epoxy glue.

After the beads were filled into the columns, the chip was mounted on the instrument and all tubing was connected, a ∼2 cm long fused silica capillary with 50 μm inner diameter and 150 μm outer diameter (TSP-050150, BGB) was cut, inserted inside the chip at the dedicated slot and glued with two component epoxy glue. Special care has been taken to avoid introducing air bubbles inside the chip.

### Blotless cryo-EM grid preparation

The blotless protein deposition on cryo-EM grid was achieved by positioning the capillary tip about 10 µm away from the grid as was estimated using a digital microscope (Dino-Lite AM9715MZTL, RS, Cat. Nr.: 216-7385). A volume between 40 and 100 nl was dispensed on the EM grid followed by suction through the same capillary by applying negative pressure (-20 mbar) through a Push-Pull pressure controller for 5-10 seconds followed by immediate grid plunging into liquid ethane.

### Blotted preparation of the grids

For experiments in which grids were blotted with dental paper points, about 100 nl of protein was deposited onto each EM grid. Dental paper points (size 40, MediStock, dental points SteriBlue) were used to manually wipe off the deposited protein in a high humidity environment for around 7 seconds and immediately plunged the grids in liquid ethane. The blotting reproducibility was improved by replacing manual with automated blotting. The microfluidic chip and blotting arm were enclosed inside a chamber, maintaining 100% humidity during blotting. Blotting with Whatman paper 4 for 0.5 seconds using a blotting angle of 8 degrees (rotation angle of the servo motor coupled to the blotting arm) yielded optimum ice thickness over many squares (see **Supplementary** Figures 4**, 9, 13**).

### Measurements of fluorescent detection sensitivity

The detection limit of proteins by fluorescence and linearity of the signal in MISO setup was measured using GFPuv (Mw 27.12 kDa). To calibrate the experimental set-up, GFP was pumped at concentrations of 0.1 mg/ml (3.7 μM), 0.01 mg/ml (370 nM) and 0.001 mg/ml (37 nM) into the detection zone of the chip at 1 μl/min flow rate (care has been taken to minimize dispersion by pumping GFP and buffer in different tubing). The fluorophore was excited at 473 nm and emitted fluorescence signal was detected at 503 nm. Volumes of 2 μl of GFP and 2 μl of buffer were alternatively pumped for six cycles for each concentration of GFP. At a concentration of 0.001 mg/ml the fluorescence signal was clearly detectable, whereas at lower concentrations it was ambiguous.

### Characterization of size-exclusion column

For SEC column characterization, a 2-column microfluidic chip with an estimated volume of 1 and 10 μl was used. The affinity column was left empty, and the SEC column was filled with Superdex 200 Increase resin (GE Healthcare). The chip was equilibrated with 20 mM HEPES pH 7.5, 150 mM NaCl, 1 mM TCEP, and 0.01% LMNG. The optical filter setup comprised GFP excitation, emission filters and GFP dichroic mirror (Thor Labs,Cat Nr: MDF-GFP).

Bio-Rad Gel filtration protein standards were prepared by diluting the protein mixture to 10 mg/ml in H_2_0. 40 μl of PBS was added to 10 μl protein, and the pH was raised by adding 5 μl NaCO_3_. 1 μl Alexa 488 fluorophore (2450 μM in DMSO) was added and incubated for 1 hour at room temperature. The excess dye was removed by ZEBA spin-columns pre-equilibrated with PBS. A volume of 20 μl labeled protein mixture was loaded onto the chip through the empty column from which 0.5 μl was diverted onto the size exclusion column at a flow rate of 1 μl/min. 1.5 μl fractions were manually collected, 1 μl was loaded onto SDS-PAGE and the unstained gel was visualized using 488 nm in Odyssey M (Li-COR).

The plate number was estimated as:

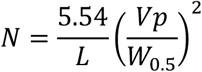

where N is number of plates per meter, L – column length (m), Vp – elution volume of the peak, W_0.5_ – peak width at half height.

### Fluorescent labeling of proteins

A fluorescent dye Chromeo P503 (Sigma-Aldrich) containing pyrylium group, which reacts with primary amines to form fluorescent adduct, was used in this study. The dye becomes fluorescent only after covalently binding to amino acids and its emission is spectrally well separated from the excitation [57]. One milligram of Chromeo P503 (Signm-Aldrich) was dissolved in 50 μl of dimethylformamide (DMF), aliquoted into 20 µl fractions and stored at -20°C. For labeling, protein extract was mixed with the dye solution in ratios 20:1 to 10000:1, for most experiments the ratio was 50:1 by volume, incubated for 1 h on ice, and loaded onto the microfluidic chip.

### Expression of β-galactosidase in E. coli

Cytoplasmic extract from *E. coli* expressing β-galactosidase [42] was prepared either by standard method growing large cultures or collecting cells directly from a Petri dish. In the first case, 12 l of *E. coli* culture were grown at 37°C in LB media supplemented with 50 µg/ml kanamycin until OD_600_ reached 0.6. The expression was induced with 1 mM IPTG for 4 hours at 37° C and the pellet, after centrifugation, was mixed with 400 ml of ice-cold lysis buffer (50 mM Tris pH 8.0, 200 mM NaCl, 2 mM MgCl_2_, 0.1 mg/ml lysozyme, DNase and inhibitors). After lysis, the suspension was centrifuged for 30 minutes at 39,000 x g (JA20, Beckman Coulter) filtered through a 0.22 μm syringe filter, aliquots were flash-frozen in LN and stored at -80°C. In the second case, one cell colony was scrapped from an LB-agar plate, supplemented with 50 µg/ml kanamycin, the cells were suspended in 1 ml of LB media with 50 µg/ml kanamycin and 1mM IPTG and incubated for 4 h at 37° C. OD_600_ of 0.85 was measured using nanodrop (NanoDrop One, ThermoFisher Scientific). The cells were centrifuged at 20,000 x g and the resulting pellet was resuspended in 30 μl of the lysis buffer. This suspension was centrifuged for 30 minutes at 20,800 x g at 4° C and 40 μl of supernatant was collected. 35 μl of this supernatant was used for purification.

### Construct design and generation of stable cell lines for btTMEM206 and mTMEM16F

The *Bos taurus* homologue of the proton-activated chloride channel TMEM206/PACC1 (btTMEM206, UniprotID: Q2KHV2) and the mouse homologue of the lipid scramblase TMEM16F (Uniprot: Q6P9J9), identified as well-expressing homologues in a small-scale expression screen. BtTMEM206 was cloned from the IMAGE clone IRCJp5010B0531D (Source BioSciences) and mTMEM16F was synthesized (Genscript) and cloned into a modified pcDNA5/FRT/TR expression vector (Invitrogen/Thermo Scientific). For this, the entire expression cassette of pcDXC3GMS (Addgene #49031) with a C-terminal Venus YFP (flanked by the KpnI sites) and a streptavidin-binding protein tag (SBP-tag) was excised with HindIII and ApaI and ligated into HindIII/ApaI-digested pcDNA5/FRT/TR to generate p5TO-CV. Subsequently, SapI-flanked PCR products of *bt*TMEM206 and *m*TMEM16F were ligated into p5TO-CV using FX cloning [58]. Stable, tetracycline-inducible cell lines were generated by co-transfection of Flp-In T-REx293 cells with pOG44 and p5TO-CV-btTMEM206 or p5TO-CV-mTMEM16F, respectively, according to the manufacturer’s instructions (Flp-In T-REx293 cell manual, Invitrogen/Thermo Scientific) and subsequent selection at a hygromycin-B concentration of 100 µg/ml. Single colonies were picked and grown for four passages in 10 cm-dishes under antibiotic pressure and the best expressing clones were identified by screening of purified proteins on an Agilent 1260 HPLC by SEC with a Superdex200 increase 5/150 column (Cytiva) and SDS-PAGE.

For conventional purification, a cell line without Venus-YFP at the C-terminus of *bt*TMEM206 was used. The expression plasmid p5TO-C was generated as described above for the Venus YFP tagged plasmid but instead, the expression cassette of pcDXC3MS (Addgene #49030) was shuttled into pcDNA5/FRT/TR. The btTMEM206 ORF was cloned into p5TO-C and sequence-verified before transfection, and selection was made as described for the C-terminally Venus-YFP tagged protein.

### Construct design, virus generation and expression of TRPC6

A construct was designed by cloning a gene of interest (GOI) into a transfer vector compatible with *flash*BAC^TM^ vector (Oxford Expression Technologies). Sf9 insect cells were maintained at mid-log phase (2-3x10^6^ cells/ml). For a single transfection, 300 μl cells were prepared by diluting the stock in Sf900III media (ThermoFisher, Cat. No. 12658019) to a concentration of 0.7-0.8x10^6^ cells. 1.8 μg of polyethyleneimine (PEI) was added to 200 μl of TC100 medium (ThermoFisher, Cat. No. 13055025) and mixed. To this mixture, 40 ng of *flash*BAC DNA and 200 ng of construct DNA were added, mixed and the mixture added to the Sf9 cells. After incubating the plate for 4 h at 27 °C, 400 μl of Sf900III and 100 μl of FCS (fetal calf serum) were added to the culture and further incubated at 27 °C for 5 days resulting in V0 virus. 100 μl of V0 virus was used to infect 3 ml of Sf9 insect cells (1x10^6^ cells/ml) and incubated at 27 °C for 48 h to generate the V1 virus stock.

For the generation of V2 virus, 50 ml of Sf9 insect cells (1.8x10^6^ cells per ml) were infected with 100 μl of V1 virus. A similar step was performed to generate V3 virus. For protein expression, 30 ml of V3 virus was used to transduce each liter of HEK-Expi cells (viable count of 1.6x10^6^ cells per ml). 18 h post transduction, 2 mM of sodium butyrate was added to the culture and expression was continued for 48 h. Cultures were harvested by centrifuging at 3500 rpm for 15 mins and the supernatant separated and discarded aseptically. The pellet was resuspended in PBS (1 mg wet weight of pellet in 1 ml of PBS) and stored at -80 °C. In conventional purification methods, about 2 mg of TRPC6 protein can be purified out of 10 g of cell pellet.

### MISO purification and characterization of β-galactosidase

The optical filter setup comprised the excitation (469 nm), emission (593 nm) filters and a dichroic mirror with reflection (350-540 nm) and transmission (570-950 nm) bands (FF562-Di03-25x36, Thor Labs). For all the MISO purification experiments, the MISO chip and protein extracts were cooled to 4°C. For experiments with 20 μg of starting amount of β-galactosidase, a MISO chip with two columns of 0.5 μl for affinity and 5 μl for size-exclusion was used (**Supplementary** Figure 2a**, b**). The affinity column was filled with PureCube NiNTA agarose beads (Cube Biotech filtered through 70 μm cut off filter (EASYstrainer, Greiner Bio-One)) and the size exclusion column was filled with highly crosslinked agarose Superose 6 beads (Cytiva). Prior to loading the cell lysate onto the affinity column, the affinity column in the chip was equilibrated with 35 μl wash buffer (25 mM HEPES pH 8.0, 100 mM NaCl, 20 mM imidazole and 1mM TCEP) and the size exclusion column was equilibrated with 55 μl of size exclusion buffer (25 mM HEPES pH 8.0, 100 mM NaCl, 1 mM TCEP). Cell lysate was fluorescently labeled with Chromeo P503. A volume of 1 μl of the dye (20 mg/ml in DMF) was mixed with 50 μl of cell lysate and incubated for 1 h. Next, 20 μl of this mixture was loaded onto the affinity column and washed with 120 μl of wash buffer. Elution buffer (25 mM HEPES pH 8.0, 100 mM NaCl, 500 mM imidazole and 1 mM TCEP) was already brought onto the chip through elution inlet IN3 (**Supplementary** Figure 2b) prior to loading the sample onto the chip. Before elution, 1 μl of elution buffer was pumped into the waste to eliminate possible contamination of elution buffer with wash buffer, then the elution buffer was diverted into the affinity column at a flow rate of 0.25 μl/min. The affinity elution signal was monitored at the first detection zone (DZ1) and when the elution peak was detected, 0.5 μl of the elution was diverted into the size-exclusion column. After this, size exclusion elution buffer was pumped through the size-exclusion column at 0.25 μl/min. The size-exclusion elution signal was monitored at the second detection zone (DZ2) and was either collected for characterization by SDS-PAGE or diverted onto EM-grid for negative staining or for cryo-plunging. To visualize eluted protein purity on SDS-PAGE, 1 μl fractions were collected, mixed with 5 μl of loading buffer and loaded on 5-15% gradient SDS-PAGE (Invitrogen) and ran at 170 V, 400 mA, 40 min followed by silver staining.

Β-galactosidase from a single *E. coli* colony or 1 μg -containing cytoplasmic extract was purified on a two-column chip with 0.5 μl affinity (Ni-NTA) and, 5 μl size-exclusion (Superose 6) columns. The on-chip columns were filled and equilibrated as described in the previous paragraph except for addition of 0.2% of DDM to the buffers. Cell extract obtained from the single *E. coli* colony (35 μl) was mixed with 1 μl of Chromeo P503 (20 mg/ml in DMF) and incubated for 1 h on ice. The labeled extract (25 μl) was loaded onto the affinity column and washed with 100 μl of wash buffer. The elution buffer (25 mM HEPES pH 8.0, 100 mM NaCl, 500 mM imidazole, 1 mM TCEP and 0.2 % DDM) was injected onto the chip through the same inlet as the wash buffer. When the peak in DZ1 was detected, 0.5 μl of the elution was diverted into the SEC column and eluted with elution buffer at a flow rate of 1 μl/min while monitoring fluorescence signal in DZ2. The eluate was collected in 2 μl fractions via capillary and used for SDS-PAGE as described above.

A similar experiment for cryo plunging of EM grids was peformed using a two-column chip (0.5/5 μl columns). The columns were packed and equilibrated as described for the previous experiment. Prior to loading the cell lysate onto the affinity column, 10 μl of *E. coli* cytoplasmic extract containing around 10 μg of β-galactosidse was incubated with 0.2 μl of chromeo P503 dye for 1 h on ice. The labelled cytoplamsic extract was diluted with 40 μl of size exclusion buffer without DDM. Next, 5 μl of the diluted cytoplamsic extract containing about 1 μg of β-galactosidse was loaded onto the first column, washed with 80 μl of wash buffer and eluted as described above with 25 mM HEPES pH 8.0, 100 mM NaCl, 500 mM imidazole, 1 mM TCEP and 0.2% DDM at a flow rate of 0.25 μl/min. Fluorescence signal was monitored at the DZ1. When the elution peak was detected, 1 μl of the eluate was diverted into the size-exclusion column and eluted at flow rate of 0.25 μl/min while monitoring fluorescence at DZ2. The eluted protein was diverted onto the capillary for plunging.

### TMEM206 purification using conventional approach

150 dishes (10 cm diameter) at a confluency of 60% were induced for protein expression with 3 μg/ml tetracycline for four days. Cells were collected from the dishes, harvested by centrifugation (4000 rpm for 15min) and frozen in liquid nitrogen. For purification, all steps were carried out at 4 °C. Frozen cells were thawed and extracted with a buffer containing 50 mM HEPES-NaOH pH 7.5, 250 mM NaCl, 10% glycerol, protease inhibitors (cOmplete, Roche) and 2% Glycodiosgenin (GDN, Anatrace/ Molecular Dimensions) for 1 hour at 4°C. Unsolubilized protein and cell debris were removed by ultracentrifugation at 41,000 rpm for 30 min using a 45Ti rotor (Beckman). The supernatant was incubated with streptavidin resin (Pierce UltraLink) for 1.5 hours on a rotary device. Unbound protein was washed off using a buffer containing 30 mM Hepes-NaOH pH 7.5, 150 mM NaCl, 8% glycerol and 0.0063% GDN and eluted with wash buffer containing 5 mM biotin in addition. The eluted protein was concentrated to 5.5 mg/ml in a volume of 100 μl and frozen in liquid nitrogen. Next, 550 μg of protein (the amount purified from 150 dishes) was thawed on ice and injected into a Superdex 200 column (5/150) equilibrated with 10 mM HEPES-NaOH pH 7.5, 150 mM NaCl, 0.063% GDN on an Agilent 1260 HPLC. The peak fraction with protein at a concentration of 1.3 mg/ml was immediately used for plunging.

### TMEM206-YFP and TMEM16F-YFP expression and solubilization for MISO

For expression of *bt*TMEM206-YFP and *m*TMEM16F-YFP, a 10 cm dish at a confluency of 80% was induced by addition of 3 µg/ml tetracycline. 24 hours later, 8 mM Na-butyrate was added, and the cells were collected after two consecutive days of expression and aliquoted in portions of half a dish per vial for *bt*TMEM206-YFP and one dish per vial for *m*TMEM16F-YFP. The pellets (∼20-30 µl volume) were frozen in liquid nitrogen and stored at -80°C.

For solubilization of *bt*TMEM206-YFP and *m*TMEM16F-YFP, a pellet of cells corresponding to half and one 10 cm-dish, respectively, was lysed in 150 µl of solubilization buffer (200 mM NaCl, 25 mM HEPES-Na, pH 7.6, 8% glycerol, 2% glyco-diosgenin (GDN) containing protease inhibitors (cOmplete, Roche)) for one hour at 4°C on a rotator. After centrifugation (20,000 g, 4°C for 30 minutes), 140 µl of the supernatant was transferred to a new tube and spun under the same conditions for another 30 minutes. Clarified supernatant (120 µl) was transferred to a new tube and spin filtered using Costar spin-X centrifuge tubes with 0.22 μm pore size at 4°C, 13,100 x g for 5 min. The filtered sample was collected (100 μl) and 80 μl was loaded onto the MISO chip.

### MISO purification and characterization of TMEM206-YFP and TMEM16F-YFP

The optical filter setup comprised excitation (469 nm) and emission (525 nm) filters and dichroic filter (reflection band 452-490nm, transmission band 505-800 nm). Pierce High-capacity streptavidin agarose resin (Thermo Fisher Scientific) was loaded into the single 3 μl column on the chip. The column was equilibrated with 80 μl of equilibration buffer (150 mM NaCl, 30 mM HEPES-Na, pH 7.6, 0.0063% GDN). Filtered solubilize (80 μl) was loaded onto the column, washed with 160 μl of equilibration buffer and eluted with elution buffer (150 mM NaCl, 30 mM HEPES-Na, pH 7.6, 0.0063% GDN, 10 mM D-(+)-biotin) at a flow rate 0.25 μl/min. The fluorescence signal was detected in DZ1 and 1 μl fractions were collected. When collecting the fractions, a delay volume of 0.7 μl between the detection zone and the tip of capillary, was accounted for. The collected fractions were used for assessing protein purity by SDS-PAGE as described above and for negative stain EM.

### MISO purification and characterization of TRPC6

The optical filter setup was identical to the one described for the previous experiment. Pierce High-capacity streptavidin agarose resin (ThermoFisher Scientific, 20357) was loaded into a 10 μl single column chip. The column was equilibrated with 100 μl of wash buffer 0.01% (150 mM HEPES pH 7.5, 150 mM NaCl, 1 mM TCEP, 0.01% LMNG and 0.0006% CHS). 100 μl of solubilizate was loaded onto the column and washed with 60 μl of 0.01% wash buffer using a flow rate of 2 μl/min. The protein was eluted using wash buffer 0.003% (same as 0.01% but with 0.003% LMNG and 0.0006% of CHS) supplemented with 15 mM D-(+)-biotin at a flowrate of 2 μl/min. During each step, the signal was measured in DZ1 and the LED intensity was set to 10. 4 μl fractions were collected and analysed using silver-stain SDS-PAGE (Pierce Silverstain kit, Thermo Fisher). The resulting gel was visualized using auto-exposure mode at wavelength of 688 nm in Odyssey M (Li-COR) to evaluate the purity of the fractions. The same experiment was repeated using a 1 μl single column chip and the undiluted fractions were analysed using negative stain.

The optimized purification was performed on a 3 μl single column chip filled with Pierce High-capacity streptavidin agarose resin. The column was equilibrated with 80 μl of wash buffer 0.01%. 80 μl spin-filtered (0.22 μm Costar SPIN-X) solubilizate was loaded onto the column and washed stepwise with 40 μl wash buffer 0.01%, 40 μl wash buffer 0.01%-ATP (buffer 0.01% supplemented with 10 mM MgCl_2_ and 10 mM ATP) and 40 μl wash buffer 0.01% at a flowrate of 2 μl/min. During loading and washing, the signal was measured in DZ1 and the LED intensity was set to level 10. The protein was eluted with wash buffer 0.003% supplemented with 15 mM D-(+)-biotin at a flowrate of 0.25 μl/min and the LED intensity reduced to the level 5. A volume of 40 – 100 nl of eluted protein was dispensed onto cryo-EM grids and plunged as described below.

For detailed biochemical characterization of TRPC6, a 2-column MISO chip was used with doubled height. 1 μl affinity column was loaded with Pierce High-capacity streptavidin agarose resin and the downstream 10 μl column was loaded with Superose 6 Increase resin (GE Healthcare). Both columns were equilibrated with 80 μl of wash buffer 0.01%. 100 μl spin-filtered (0.22 μm) solubilizate was loaded onto the affinity column and washed stepwise with 40 μl wash buffer 1%, 40 μl wash buffer 0.1%, 40 μl wash buffer 0.01%, 40 μl wash buffer 0.01%-ATP, and 40 μl wash buffer 0.01% at a flowrate of 2 μl/min to gently reduce the detergent concentration. During loading and washing of the affinity column, the signal was measured in DZ1 and the LED intensity level was set to 10. The protein was eluted using wash buffer 0.01% supplemented with 15 mM D-(+)-biotin at a flowrate of 0.25 μl/min and the LED intensity was reduced to level 5. 1 μl of eluted protein was injected into the gel filtration column and the fluorescence detection position switched DZ2 while light intensity was increased to level 10. 1 μl fractions were collected manually from the tip of the outlet capillary and used for negative stain and SDS-PAGE analysis. For negative stain EM, fractions were diluted 5 times with wash buffer 0.01% and 2 μl of diluted sample was applied on a freshly glow-discharged carbon grid (30 sec, 5 mA in air using ELMO glow discharge system (Cordouan Technologies)). For SDS-PAGE analysis, leftover fractions were diluted 4 times with loading buffer. After SDS-PAGE, the unstained gel was visualized using fluorescent imaging in Odyssey M (Li-COR) using auto-exposure and excitation wavelength of 488 nm and subsequently silver-stained. The stained gel was visualized using fluorescent imaging with excitation wavelength of 700 nm (Li-COR Odyssey M).

### Quantification of protein amounts. (Concentration of β-galactosidase in E. coli extract)

To quantify βG, the activity of *E. coli* expressing β-galactosidase cell extract was determined using the βG Assay Kit (Thermo Fisher Cat. Nr. K1455-01). Cell extracts were diluted 1000-fold in 25 mM HEPES pH 8.0, 100 mM NaCl, 1 mM TCEP buffer supplemented with 0.1% DDM. 5–40 μl diluted extracts were further diluted with the buffer to a volume of 120 μl. 15 μl OPNG and 15 μl 10x cleavage buffer were added, and the samples were incubated for 15 min at 37°C. To stop the reaction, 150 μl NaCO3 was added and the OD_410nm_ was measured against a blank sample containing OPNG and cleavage buffer. The total amount (nmoles) of cleaved OPNG molecules were determined as:

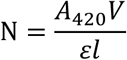

where N- of hydrolyzed ONPG (nM), A_420_ – absorbance at WL 420 nm, V- total volume in nl and ε - extinction coefficient (4500 nl nM^-1^ cm^-1^), l – optical path length (cm).

The specific activity of the lysate was determined as:

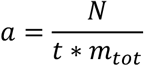

where, a - specific activity (nM min^-1^ mg^-1^), t -reaction time (min), m_tot_ - total protein amount (mg). The total protein was determined using the Pierce BS protein assay (Thermofisher). A standard β- galactosidase activity curve was determined by plotting the OD_410nm_ against different amounts of purified β-galactosidase to determine the specific activity of the enzyme. From this the concentration of β-galactosidase present in the *E. coli* cell extract used for the experiments was determined as 1.40 μg/μl.

### Conventional cryo-EM grid plunging for btTMEM206

The cryo-EM grids were plunged using a Leica CP-3 plunger. A volume of 3 μl was applied to R1.2/1.3- Cu 300 mesh grids (Quantifoil) that were glow discharged (ELMO, Cordouan Technologies) at 9 mA for 1 min, blotted from one side in CP-3 cryoplunge (Gatan) at room temperature with Whatman 1 filter paper for 2 seconds at 90% relative humidity.

### Cryo-EM grid preparation for β-galactosidase purified on MISO

For βG purified starting from 20 μg containing cell extract, volumes of 50-100 nl of the eluate from size-exclusion column were deposited on plasma cleaned (100 W, 50 kHz, 1 min in CUTE (Femto Science)) R2/1 holey carbon copper 300 mesh grids (Quantifoil), blotted manually with absorbent paper points of size 40 (MediStock, dental points SteriBlue) in high humidity atmosphere and immediately plunged into liquid ethane. Up to 8 grids were plunged per elution peak.

For βG purified starting from 1 μg-containing cell extract, 50-100 nl of the eluate from the size-exclusion column was applied per R2/1 copper mesh 300 electron microscopy grid (Quantifoil) coated with a 3 nm carbon film. The grids were blotted automatically from the front using Whatman 4 paper strip for 0.5 s in 100% humidity and immediately plunged into liquid ethane. Prior to protein deposition the grids were plasma cleaned at 25 W, 50 kHz for 15 s in CUTE.

### Cryo-EM grid preparation for TMEM206-YFP purified on MISO

Volumes of 60-100 nl of *bt*TMEM206-YFP eluted from the on-chip column through a capillary were deposited onto carbon R2/1, R1.2/1.3 and UltraAUFoil R1.2/1.3 grids. The elution peak between volumes of 2.3 and 4.4 μl was fractionated. The grids were blotted with Whatman 4 filter paper for 0.5 s in 100% humidity chamber before plunging. The gold UltraAUFoil R1.2/1.3 grid that was plunged close to the peak had the highest density of particles and was used for cryo-EM data collection.

### Cryo-EM grid preparation for TMEM16F-YFP purified on MISO

The eluted protein was plunged on holey and gold UltraAUFoil R1.2/1.3, 300 mesh grids alternatively from 3.7-4.6 - μl. The grids were deposited with about 60-100 nl of protein from the capillary and were blotted with Whatman 4 filter paper for 0.5 s in 100% humidity chamber. The gold UltraAUFoil R1.2/1.3 grid that was plunged close to the peak at 4.5 μl had the highest density of particles and was used for further collection and processing.

### Cryo-EM grid preparation for TRPC6 protein purified on MISO

During each plunging session, different grid types were used to sample a range of plunging conditions. Quantifoil carbon R1.2/1.3 Cu300 were glow-discharged for 1 min at 60 W, gold UltrAUFoil R1.2/1.3 grids were glow-discharged for 5 min at 100 W using CUTE. Protein eluted in the volume between 3.3 and 4.8 μl was plunged by applying 60–100 nl onto the freshly glow-discharged grids and front blotting for 0.5 sec using Whatman 4 filter paper. The blotting angle was 8 degrees, and the humidity was 100%. Grids were screened on the JOEL CryoARM300 electron microscope and an UltraAUFoil R1.2/1.3 grid displaying best contrast and particle density was used for data collection. As a control, 50 nl fractions were collected in between plunged grids and after diluting around 5x with wash 0.01% buffer used for negative stain analysis.

### Cryo-EM data collection

All cryo-EM datasets were collected on JEOL CryoARM300 transmission electron microscope (TEM) equipped with an Omega Filter (operated with slit centered on zero loss peak with width set to 20 eV slit) using SerialEM v3.8.0 [7]. Micrographs (exposure time 2.796 s) were recorded on a Gatan Summit K3 direct electron detector operating in counting mode with each movie fractionated into 61 frames and total dose of around 60 e^-^/Å^2^ at a nominal magnification of 60,000, a nominal magnified pixel size of 0.7-0.76 Å and dose rate in the range 9.1-11.5 e^-^/sec pixels.

The *E. coli* β-galactosidase dataset collected from grids prepared from 20 and 1 μg containing cell extract comprised of 4,965 and 6,055 movies collected using a defocus range of 0.8-1.7 μm and 1-1.8 μm, respectively. For cryo-EM dataset collected from *bt*TMEM206 purified and plunged by conventional approach, 6,398 images were collected using a 3×3 multi-shot pattern, with 27 exposures per position and a defocus range of 0.7-1.7 μm. The *bt*TMEM206-YFP and mTMEM16F-YFP dataset collected from grids plunged using MISO device were collected using a pattern of 3×3 multi-shot pattern, comprised 18,801 and 15,039 movies recorded using a defocus range of 1-1.8 and 1-1.7 μm, respectively, using 5×5 multi-shot pattern and 3 exposures per hole. The dataset collected from TRPC6 comprised 5,573 movies collected with defocus range of 0.8-2.8 μm using a 5×5 multi-shot pattern and 3 exposures per hole. Each movie consisted of 61 frames and was recorded using a flux of 18.2 e-/sec pixels.

### Cryo-EM Data processing of β-galactosidase from 20 ug

The movies were preprocessed in CryoSPARC Live v4.4.1 for motion correction using Patch Motion correction and Patch CTF was used to estimate the contrast transfer function (CTF) of the motion-corrected images. All further steps were completed in cryoSPARC v4.5.3. Micrographs containing ice contaminants were sorted out resulting in 3994 clean micrographs. Blob picking followed by inspection has yielded 1,303,487 particles that were extracted into box of 392 pixels Fourier cropped to 64 pixels. Following 2D classification, 1,289,528 particles were selected and used to generate *ab initio* 3D volume reconstruction with 4 classes followed by several rounds of heterogenous refinements using the final reconstructions from each round as an input map for the next refinement (**Supplementary** Figure 6). 739,896 particles from the best resolved 3D class were selected and re-extracted into boxes of 392 pixels Fourier cropped to 196 pixels and *ab initio* reconstructions followed by heterogeneous refinement were repeated followed by selection of particles from the best resolved 3D class. For the final refinement, 599,438 particles were selected and extracted into box of 392 pixels Fourier cropped to 276 pixels. The 3D map was further refined using homogeneous refinement, local per particle CTF refinement and non-uniform refinement to yield a map at 2.3 Å resolution reconstruction.

### Cryo-EM data processing of β-galactosidase from 1 ug

Initial processing steps were identical to the processing described above. A total of 6,055 micrographs were collected and after removing micrographs contaminated with ice 5,734 micrographs were retained. Topaz particle picking followed by particle extraction into 400 pixels box Fourier cropped to 64 pixels yielded 238,014 particles that were subjected to one round of 2D classification after which 142,129 particles were selected. *Ab initio* reconstruction with 3 classes was calculated followed by, heterogenous refinement after which one 3D class containing 81,127 particles was retained. The selected particles were re-extracted in the box size 400 pixels Fourier cropped to 276 pixels and 3D maps were refined by homogeneous and non-homogeneous refinements. Next, particles were re-extracted into a box of 400 pixels without decimation (a total of 80,443) followed by homogeneous and non-uniform refinements, ctf refinement, motion correction and homogeneous refinement. Resulting reconstruction had resolution of 2.16 Å to which sharpening B-factor of - 49.4 Å^2^ was applied to produce final 3D map (**Supplementary** Figure 7).

### Cryo-EM data processing for conventional btTMEM206 preparation

Micrographs were pre-processed in CryoSPARC Live v4.4.1 as described above. Subsequently, all steps were done in cryoSPARC v4.5.3. Images with ice contamination were removed resulting in 5,735 clean micrographs from which 24,838,674 particles were picked using a multi-picking approach (blob, topaz and template picker) and extracted into a box of 400 pixels Fourier cropped to 100 pixels. Particles and templates resulting from a small subset from blob picking were used as inputs for template and topaz picking, respectively. Several rounds of 2D classification were conducted and classes containing 7,960,000 particles were selected. The resulting particle stack was reconstructed *ab initio* with four volumes after which 2,429,256 particles from the best reconstruction were used for another *ab initio* reconstruction of two volumes. 1,671,826 particles from the best class were further cleaned through several heterogenous refinements using two *ab initio* maps as initial model. After removal of duplicates 332,041 particles were retained and re-extracted into a box of 400 pixels. Following non-uniform refinement, a map at resolution of 2.86 Å was calculated and sharpened by applying a B-factor of -109.9 Å^2^ (**Supplementary** Figure 8).

### Cryo-EM data processing of mTMEM206-YFP prepared using MISO

Micrographs were pre-processed in CryoSPARC Live v4.4.1 as described above. Subsequently, all steps were done in cryoSPARC. After image selection 7,366 micrographs were retained and 668,604 images picked using template picker. The images were extracted into boxes of 400 pixels Fourier cropped to 100 pixels followed by 2D classification. 633,520 particles were selected and used for *ab initio* reconstruction with 5 classes followed by hetero refinement. One class displaying features of TMEM206 and containing 101,757 particles was selected and particles were re-extracted into a box of 400 pixels Fourier cropped to 200 pixels. Next, non-uniform refinement followed by global CTF refinement and another round of non-uniform refinement was performed. After 3D classification into 6 classes 87,106 particles were selected and reconstructed by non-uniform refinement to resolution of 3.0 Å to which a sharpening B-factor of -103.6 Å^2^ was applied (**Supplementary** Figure 10).

### Cryo-EM data processing of mTMEM16F-YFP prepared using MISO

The movies were preprocessed in CryoSPARC Live v4.4.1 for motion correction using Patch Motion correction and CryoSPARC CTF estimation was used to estimate the contrast transfer function (CTF) of the motion-corrected images. All further steps were accomplished in CryoSPARC. Micrographs containing ice contaminants were sorted out resulting in 12,521 clean micrographs. A multi-picking approach combined blob, topaz and template picking and yielded 6,478,121 particles that were subjected to several rounds of 2D classification after which 2,001,905 particles were selected. The resulting particle stack was split into 2 subsets and classified into 2 and 3 classes *ab initio* followed by hetero refinements (**Supplementary** Figure 11). Particles from the two best resolved classes (a total of 402,208) were merged and extracted into a 400 pixels box without binning. Duplicates were removed to yield 184,435 particles which were further cleaned by hetero refinement. Selected particles (71,594) were used for non-uniform refinement and reference-based motion correction. The motion-corrected particles (69,117 particles remaining after reference-based motion correction) were refined by non-uniform refinement and applying C2 symmetry to produce reconstruction to resolution of 3.5 Å. A sharpening B-factor of -100.4 Å^2^ was applied to the final map.

### Cryo-EM image processing for TRPC6

For all datasets, image processing was done with CryoSPARC v4.5.3. Movie alignment was done with Patch MotionCorr and contrast transfer function (CTF) was estimated with Patch CTF. 6,686 movies were manually curated and the 4,563 good movies were used for Topaz particle picking implemented in CryoSPARC using the standard ResNet16 network and an estimated particle diameter of 80 Å. 186,934 particles were extracted in a box of 400 pixels, decimated to 100 pixels. To find additional side-views, template-based particle picking using a side-view reference, obtained from an initial processing round, was performed resulting in an additional 273,697 particles. Both particle sets were combined and 2 rounds of reference-free 2D classification were performed after which the best classes were selected accounting for 323,090 particles. Next, 3 *ab initio* models with C1 symmetry and a maximum resolution of 12 Å were generated. These models were used as references in heterogenous refinement using all particles and imposing C4 symmetry. A single class displaying the high-resolution features was retained (135,630 particles) and corresponding particles were re-extracted. After removing duplicate particles and re-extracting particles decimated to 250 pixels, 90,318 particles were non-uniform refined. To separate complete particles from cytosolic domain-only particles, 3D classification was performed with resolution limited of 6 Å and applying C4 symmetry. One class corresponding to full (11,201 particles) and 3 good classes corresponding to cytosolic domain only (75,449 particles) were used as a reference for heterogeneous refinement. The final volumes were obtained by performing global and local ctf refinements, reference-based MotionCorr and non-uniform refinement with C4 symmetry resulting in 3D reconstructions at resolution of 3.4 Å and 3.5 Å, respectively (**Supplementary** Figure 14).

### Cryo-EM model building and refinement

For the structure of *E. coli* β-galactosidase 20 μg, an initial model was built using ModelAngelo v.0.2.2 [13] by providing the final density map and the FASTA sequence of 1 subunit. The resulting monomer was manually rebuilt in Coot 0.9.5 [59] and complete model generated by applying a D2 symmetry in Phenix 1.19 [60]. Water molecules were automatically added, and the resulting model was refined in Phenix using the douse and real_space_refinement routine respectively [61]. In the latter, global minimization, rigid body fit, and local rotamer fitting were performed. 5 cycles of iterative building /refinement were completed to obtain the final model. The refined model was used as a starting model for the structure of *E. coli* β-galactosidase 1 μg and a similar building strategy was used.

For the structure of *bt*TMEM206-YFP, an initial model was built using Model Angelo v.0.2.2 by providing the density map and the protein sequence. A similar building strategy was followed, and the complete model was generated by applying C3 symmetry in Phenix 1.19. The final refined model was used as a starting model for *bt*TMEM206 conventional model building. For the structure of *m*TMEM16F-YFP, the model of *Mus musculus* TMEM16F (PDB 6P46 [62]) was used as a starting model for building. A monomer was fitted into the cryo-EM map in ChimeraX v.1.7 [63] and manually rebuilt using Coot 0.9.5. Poorly resolved parts of the initial model were truncated. The resulting model was expanded to a dimer by applying C2 symmetry. One coordinated Ca^2+^ ion was well resolved and modeled. The resulting model was refined in Phenix using real_space_refinement. Here, global minimization, rigid body fit, and local rotamer fitting were performed.

For TRPC6 the map was fitted with the structure of human TRPC6 PDB 6UZ8[64]. The fitted model was adjusted in Coot, refined in Phenix and used for comparison with the starting model.

The models were validated using MolProbity [65] and Coot. Figures were generated using UCSF Chimera and UCSF ChimeraX v1.7.

