## Supplementary information for "MISO: Microfluidic protein isolation enables single particle cryo-EM structure determination from a single cell colony"

### Supplementary Table 1 | Cryo-EM data collection, refinement and validation statistics

| | **$\beta$-gal 20 $\mu$g** | **$\beta$-gal 1 $\mu$g** | **TMEM206 conventional** | **TMEM206 MISO** | **TMEM16F MISO** | **TRPC6 MISO** |
|---|---|---|---|---|---|---|
| EMDB | **EMD-52333** | **EMD-52334** | **EMD-52344** | **EMD-52345** | **EMD-52346** | **EMD-52486 / EMD-52487** |
| PDB | **9HPL** | **9HPM** | **9HQN** | **9HQO** | **9HQP** | |
| **Data collection and processing** | | | | | | |
| Electron microscope | CryoARM 300 | | | | | |
| Electron detector | K3, CDS mode | | | | | |
| Voltage (kV) | 300 | | | | | |
| Nominal magnification | 60,000 | | | | | |
| Exposure time (s) | 2.796 | | | | | |
| Defocus range (µm) | 0.8 – 1.7 | 1 – 1.8 | 0.7-1.7 | 1 – 1.8 | 1 – 1.7 | 0.8 – 2.8 |
| Number of exposures per stage position | 45 | 45 | 27 | 75 | 75 | 75 |
| Number of frames | 59 | 59 | 59 | 59 | 59 | 59 |
| Pixel size (Å) | 0.71 | 0.7 | 0.76 | 0.695 | 0.695 | 0.7155 |
| Electron dose (e–/Å$^2$) | 60.5 | 60.5 | 60 | 60.5 | 60.5 | 60 |
| Number of micrographs used for particle selection | 3,994 | 5,734 | 5,735 | 16,597 | 12,521 | 4,563 |
| **3D reconstruction** | | | | | | |
| Initial particle images (no.) | 1,303,487 | 238,014 | 24,838,674 | 2,187,681 | 6,478,121 | 452,519 |
| Final particle images (no.) | 599,438 | 80,443 | 332,041 | 87,106 | 69,117 | 11,204/ 75,448 |
| Symmetry imposed | D2 | D2 | C3 | C3 | C2 | C4 |
| Map resolution (Å) FSC threshold | 2.34 | 2.16 | 2.86 | 2.97 | 3.51 | 3.49/3.44 |
| Sharpening B-factor (Å$^2$) | 84.5 | 49.4 | 109.9 | 107.5 | 100.2 | 80.5/117.3 |
| **Refinement** | | | | | | |
| Initial model used (PDB code) | Model_Angelo | Model_Angelo | TMEM206 MISO | 6P46 | Model_Angelo | - |
| Model resolution (Å) FSC threshold 0.5 | 2.4 | 2.3 | 3.0 | 3.2 | 3.9 | - |
| Map sharpening *B* factor (Å$^2$) | 84.5 | 49.4 | 109.9 | 107.5 | 100.2 | - |
| Model composition | | | | | | - |
| Non-hydrogen atoms | 34186 | 33848 | 6390 | 6330 | 10402 | |
| Protein residues | 3912 | 3928 | 786 | 792 | 1281 | |
| Ligands | 0 | 0 | 3 | 0 | 2 | |
| Waters | 2826 | 2341 | 30 | 0 | 0 | |
| *B* factors (Å$^2$) | | | | | | - |
| Protein | 24.76 | 41.23 | 27.70 | 46.92 | 51.85 | |
| Ligand | - | - | 47.58 | - | 25.47 | |
| Water | 28.69 | 47.36 | 9.43 | - | - | |
| R.m.s. deviations | | | | | | - |
| Bond lengths (Å) | 0.003 | 0.002 | 0.002 | 0.003 | 0.003 | |
| Bond angles (°) | 0.644 | 0.517 | 0.556 | 0.556 | 0.663 | |
| Validation | | | | | | - |
| MolProbity score | 1.77 | 1.61 | 1.51 | 1.59 | 2.12 | |
| Clashscore | 6.92 | 4.63 | 4.60 | 8.22 | 14.38 | |
| Poor rotamers (%) | 1.68 | 1.82 | 0.0 | 0.0 | 0.0 | |
| Ramachandran plot (%) | | | | | | |
| Favored | 96.62 | 97.00 | 96.03 | 97.20 | 92.77 | - |
| Allowed | 3.27 | 3.00 | 3.97 | 2.80 | 7.23 | - |
| Disallowed | 0.1 | 0.00 | 0.0 | 0.0 | 0.0 | - |

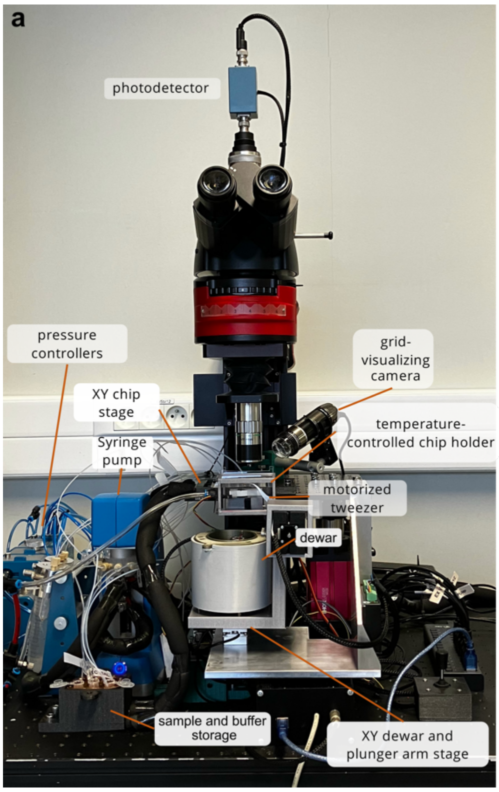


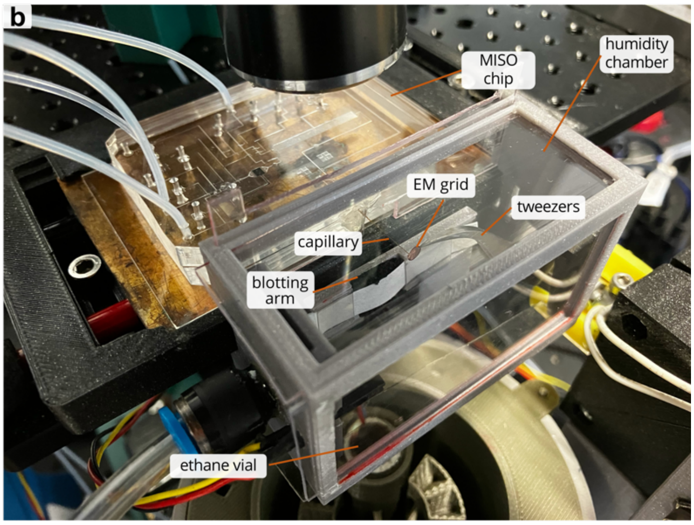


**Supplementary Figure 1 | Experimental setup. a**, An overview of the experimental setup. **b**, Closeup of the MISO chip and plunger module. Key elements are labeled.

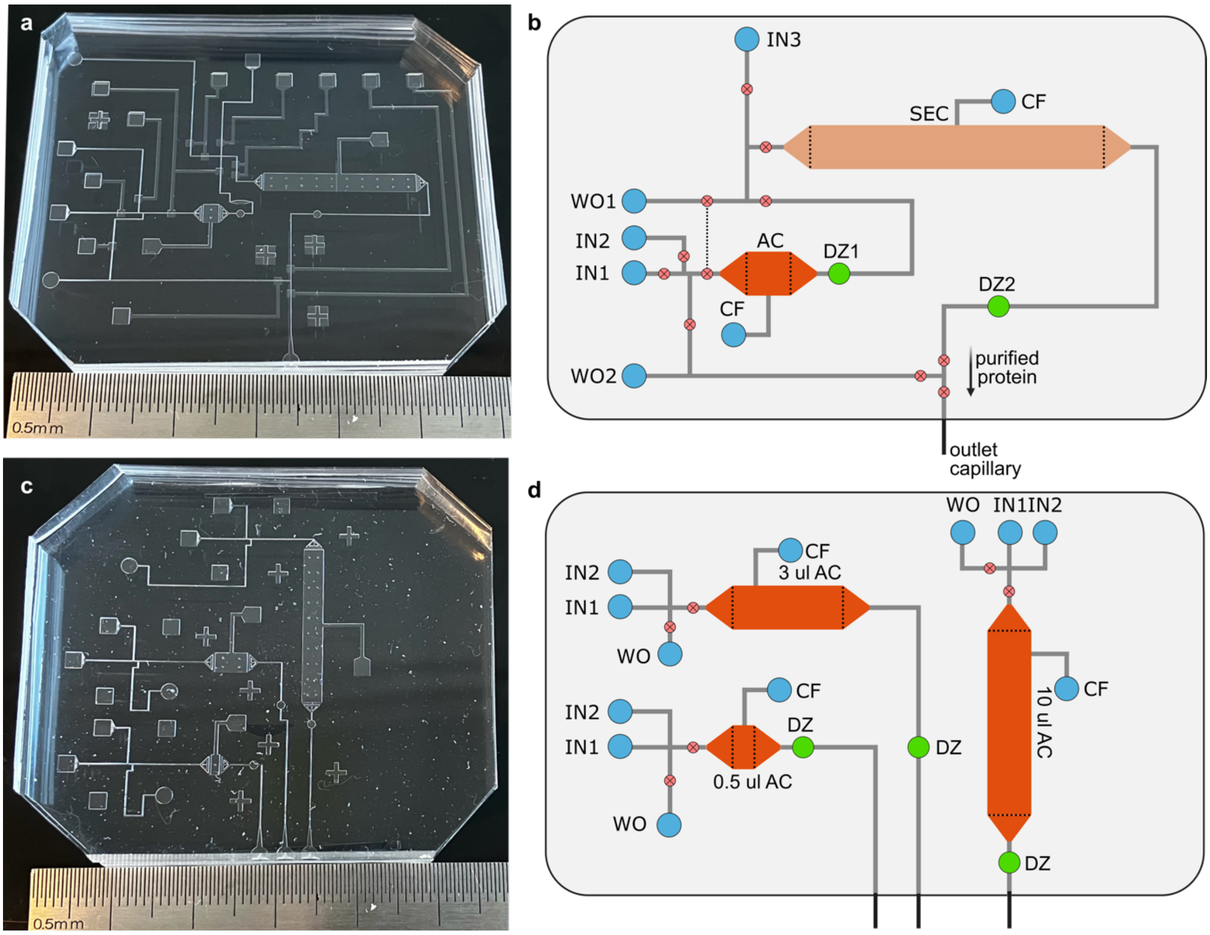


**Supplementary Figure 2 | Layout of MISO chips**. **a, b**, Chip photograph and schematics of chip elements for a 2-column MISO chip. **c, d**, Chip photograph and schematics of chip elements for a 1-column MISO chip. Three single-column microfluidics circuits with columns of 0.5, 3 and 10 μl were designed on a single chip. The valves are shown as red circles and pneumatic channels are not shown in the schematics of b and d panels. The following abbreviations are used: IN – inlet, WO – waste outlet, CF- column filling port, DZ – detection zone.

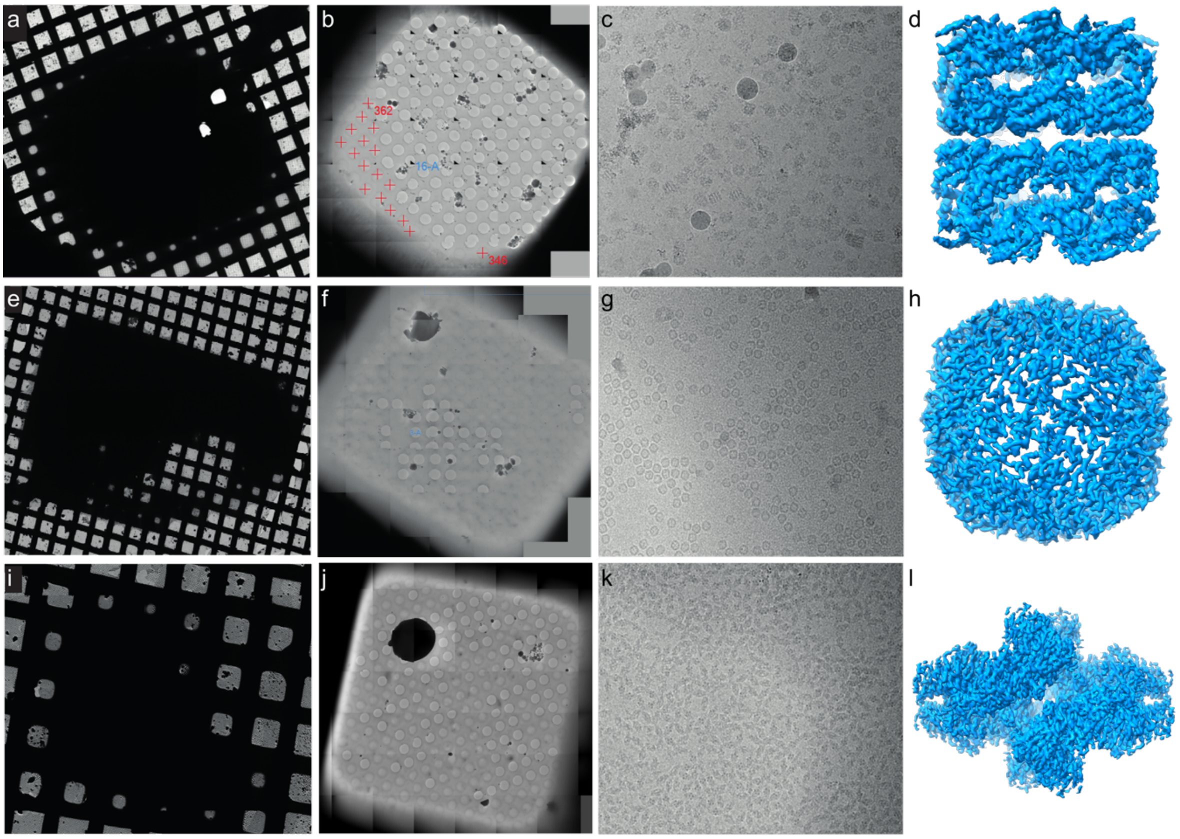


**Supplementary Figure 3 | Blotless cryo-EM grid preparation using MISO plunger.** Examples of grid atlases, grid squares, micrographs and 3D reconstructed volumes for three test proteins: *E. coli* GroEL (panels **a-d**), mouse heavy chain apoferritin (panels **e-h**) and *E. coli* β-galactosidase (panels **i-l**). The reconstructions were determined to resolution 4.0, 2.6, 2.6 Å, respectively.

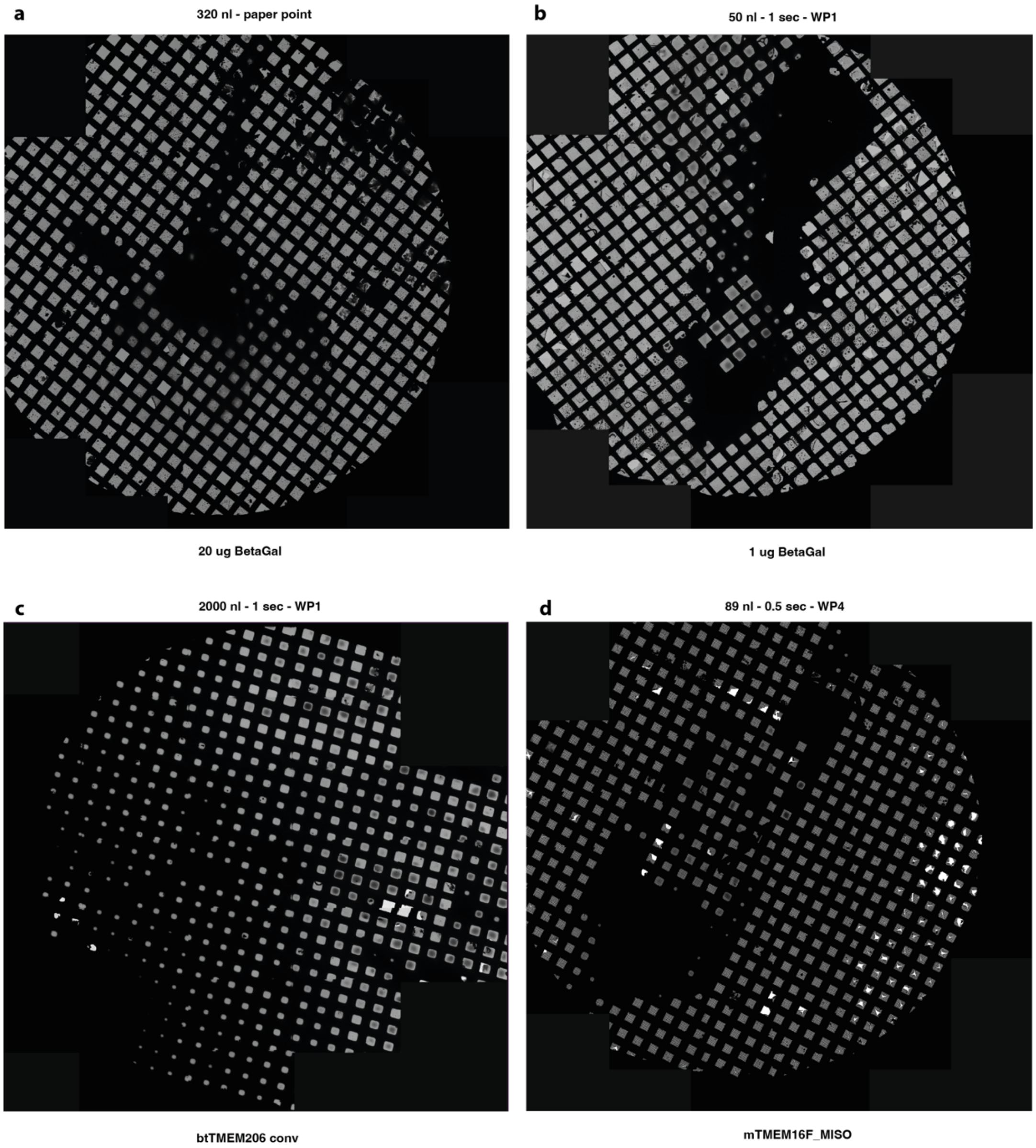


**Supplementary Figure 4 | Atlases of cryo-EM grids used for data collection. a,** 20 μg βG MISO blotted using paper point. **b,** 1 μg βG MISO plunged using Whatman paper blotting. **c,** *bt*TMEM206 purified and plunged using the conventional method. **d,** *bt*TMEM16F-YFP purified and plunged using MISO. Deposition volume and blotting conditions are indicated.

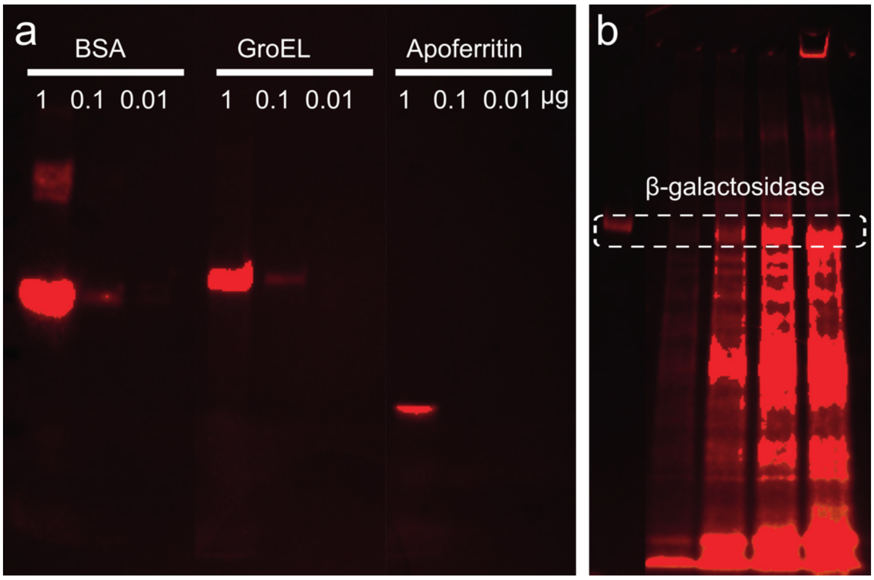


**Supplementary Figure 5 | Fluorescent protein labelling with Chromeo P503 dye. a,** SDS-PAGE images of purified bovine serum albumin, GroEL and apoferritin separated on SDS-PAGE and visualized using fluorescent imaging. The amount of protein loaded per laine is indicated in micrograms. **b**, Total cytoplasmic extract from *E. coli* expressing βG was labelled with Chromeo P503 and separated on SDS-PAGE. The total protein loaded per lane was 1 μg. The left lane is pure protein labelled with dye in the ratio of 20:1. For the lanes from 2 to 5 (left to right) the cytoplasmic extract was labelled with the dye in the ratio of 100:1, 20:1, 10:1 and 5:1.

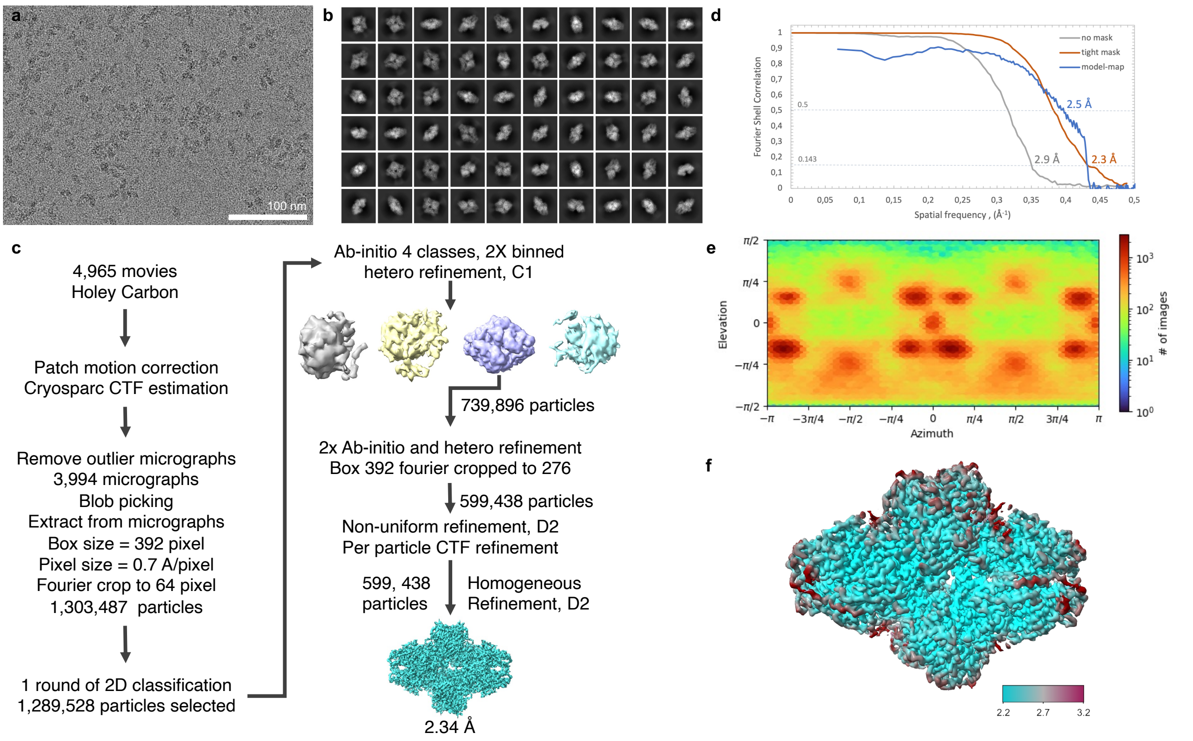


**Supplementary Figure 6 | Processing of β-galactosidase data for cryo-EM sample prepared starting from 20 μg βG containing cytoplasmic extract. a**, A micrograph, **b**, 2D class averages. The box size is 278 Å. **c**, Image processing scheme. **d**, Fourier Shell Correlation curves for unmasked and masked half-maps and between model and map. **e**, Heat map shows distribution of particle orientations. **f**, 3D map surface colored by local resolution.

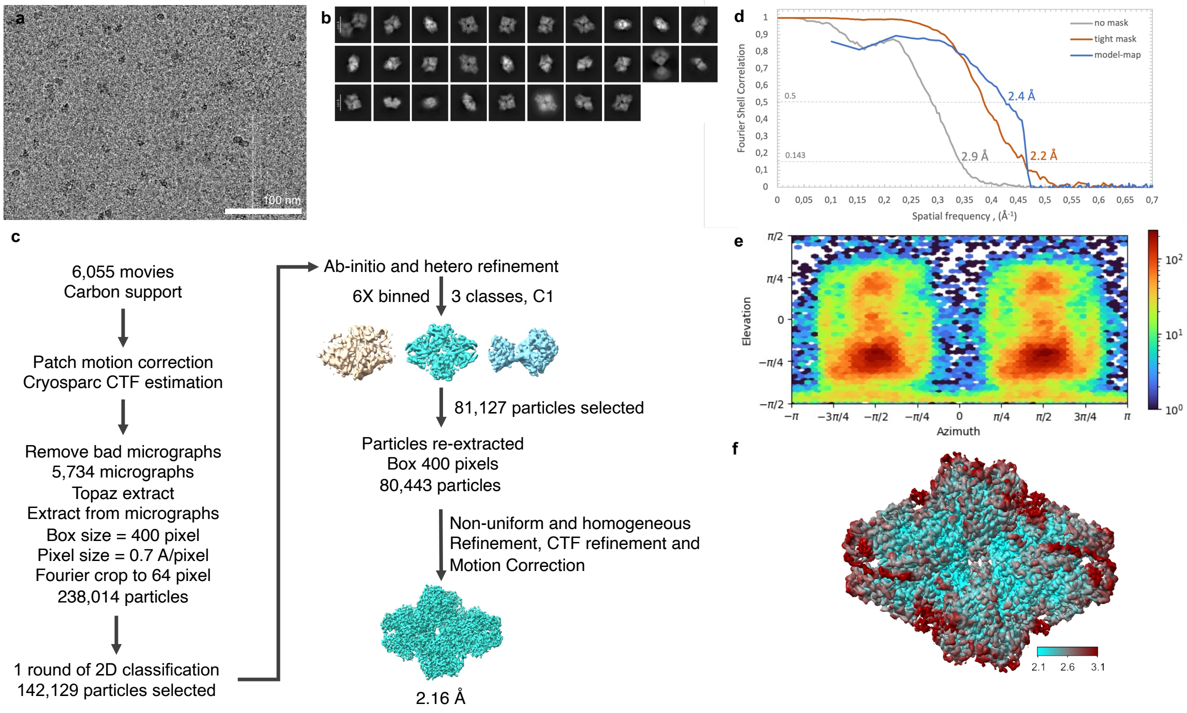


**Supplementary Figure 7 | Processing of β-galactosidase data for cryo-EM sample prepared starting from 1 µg βG containing cytoplasmic extract. a**, A micrograph, **b**, 2D class averages. The box size is 280 Å. **c**, Image processing scheme. **d**, Fourier Shell Correlation curves for unmasked and masked half-maps and between model and map. **e**, Heat map shows distribution of particle orientations. **f**, 3D map surface colored by local resolution.

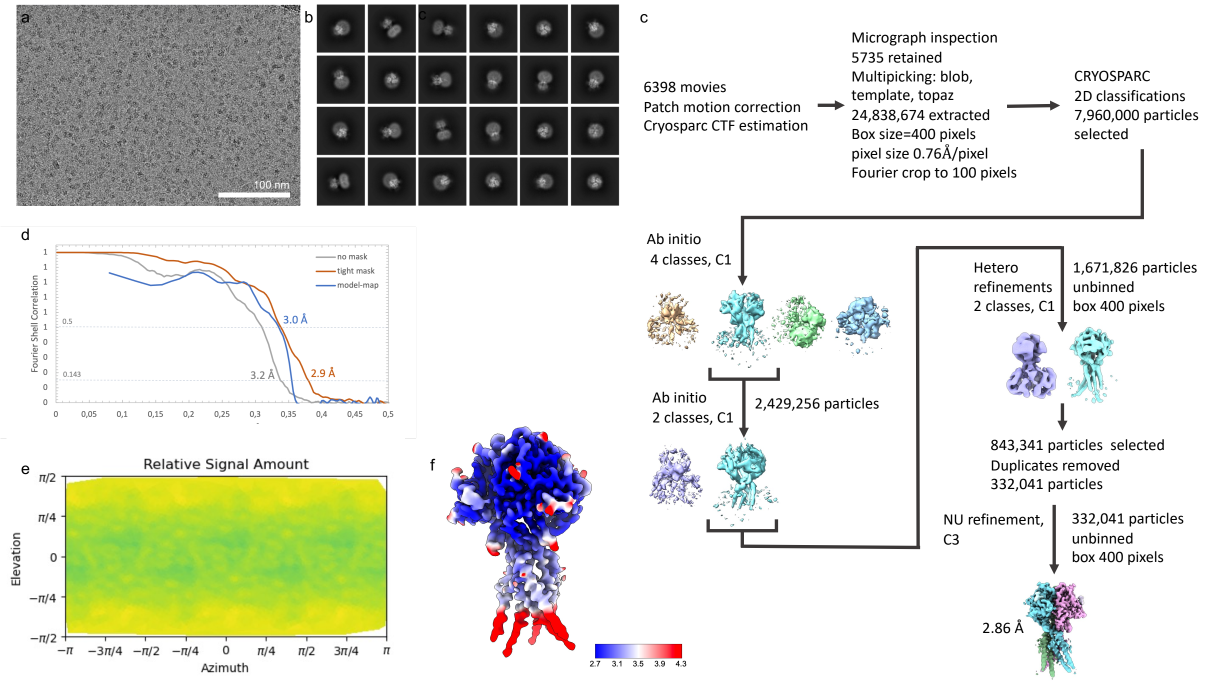


**Supplementary Figure 8 | Processing of TMEM206 purified and plunged using conventional approach. a**, A micrograph, **b**, Selected 2D class averages. **c**, Image processing scheme. **d**, Fourier Shell Correlation curves for unmasked and masked half-maps and between model and map. **e**, Heat map shows distribution of particle orientations. **f**, 3D map surface coloured by local resolution.

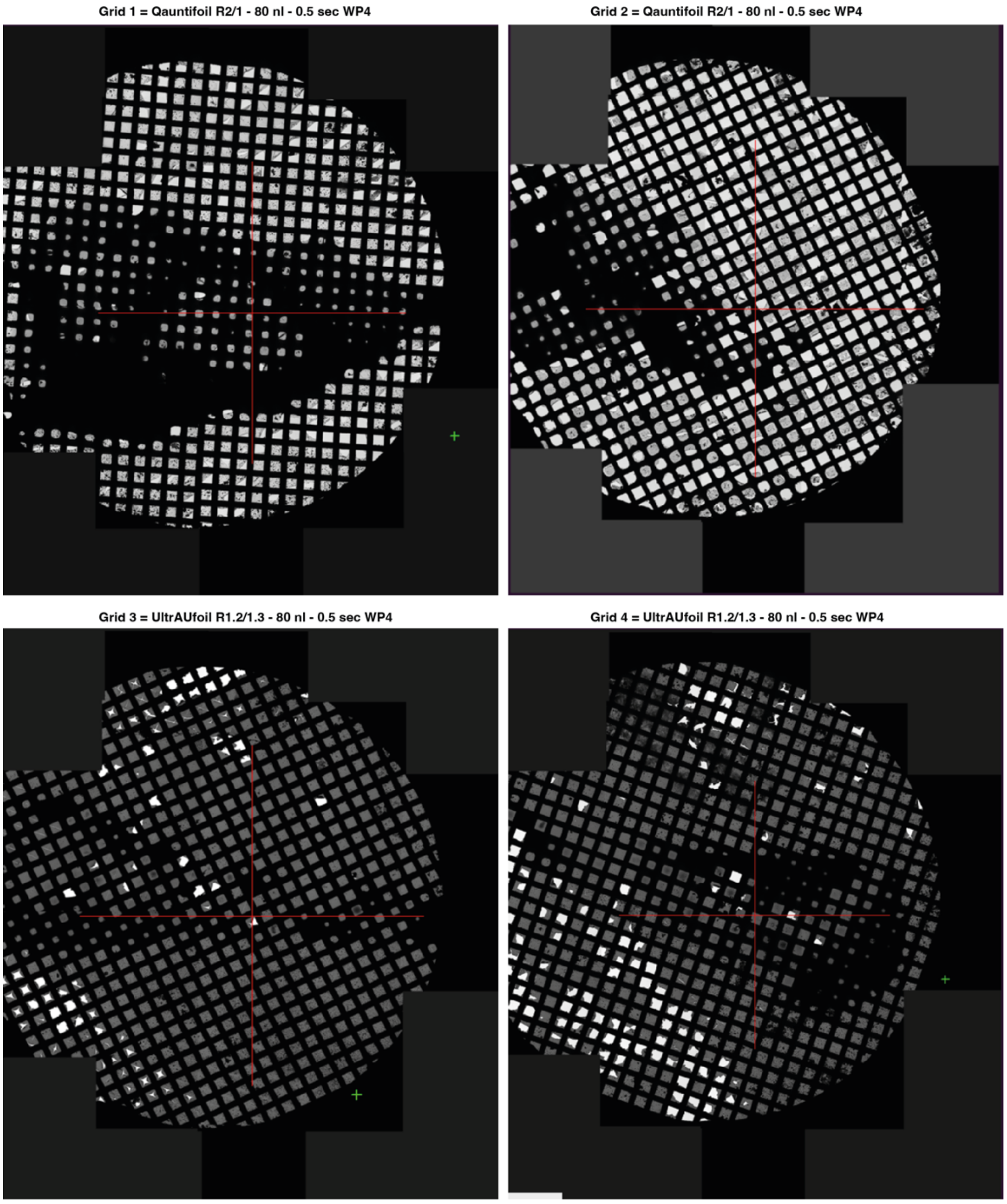


**Supplementary Figure 9 | Atlases of the *bt*TMEM206-YFP grids plunged using MISO.** The atlases are shown for 4 consecutively plunged grids. Deposited protein volume, blotting time and blotting paper type are indicated.

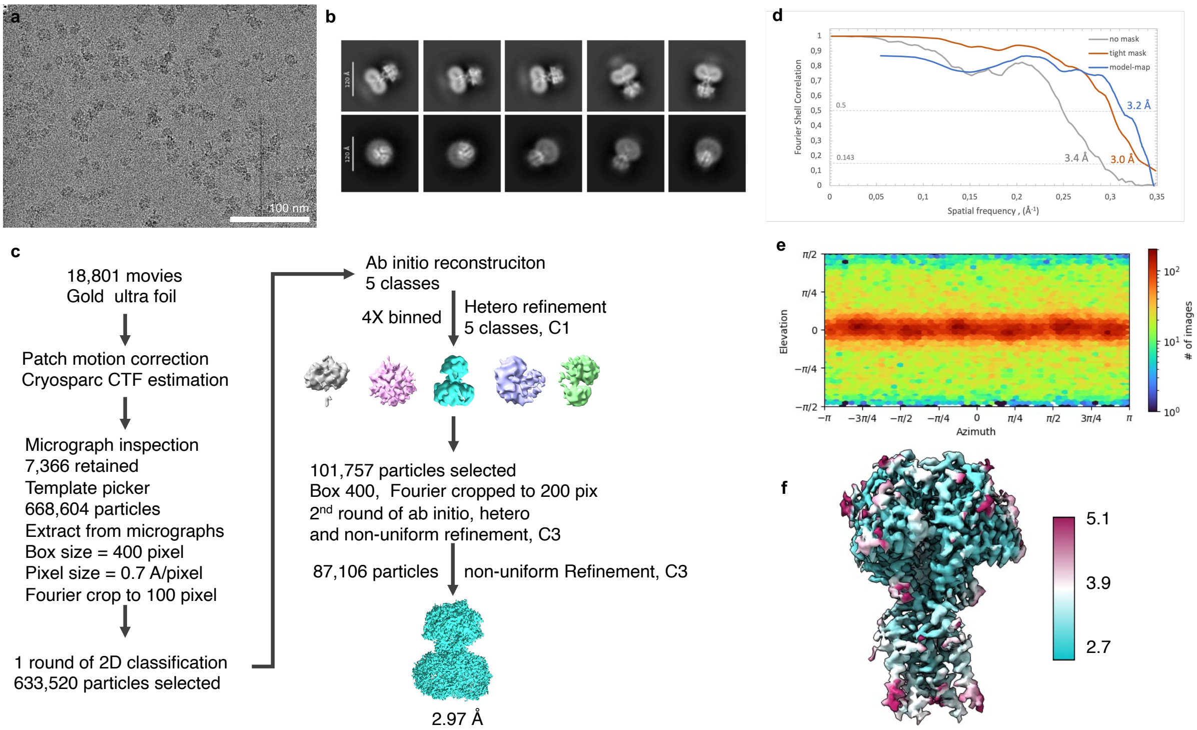


**Supplementary Figure 10 | Processing of TMEM206-YFP purified and plunged using MISO. a**, A micrograph, **b**, Selected 2D class averages. **c**, Image processing scheme. **d**, Fourier Shell Correlation curves for unmasked and masked half-maps and between model and map. **e**, Heat map shows the distribution of particle orientations. **f**, 3D map surface coloured by local resolution.

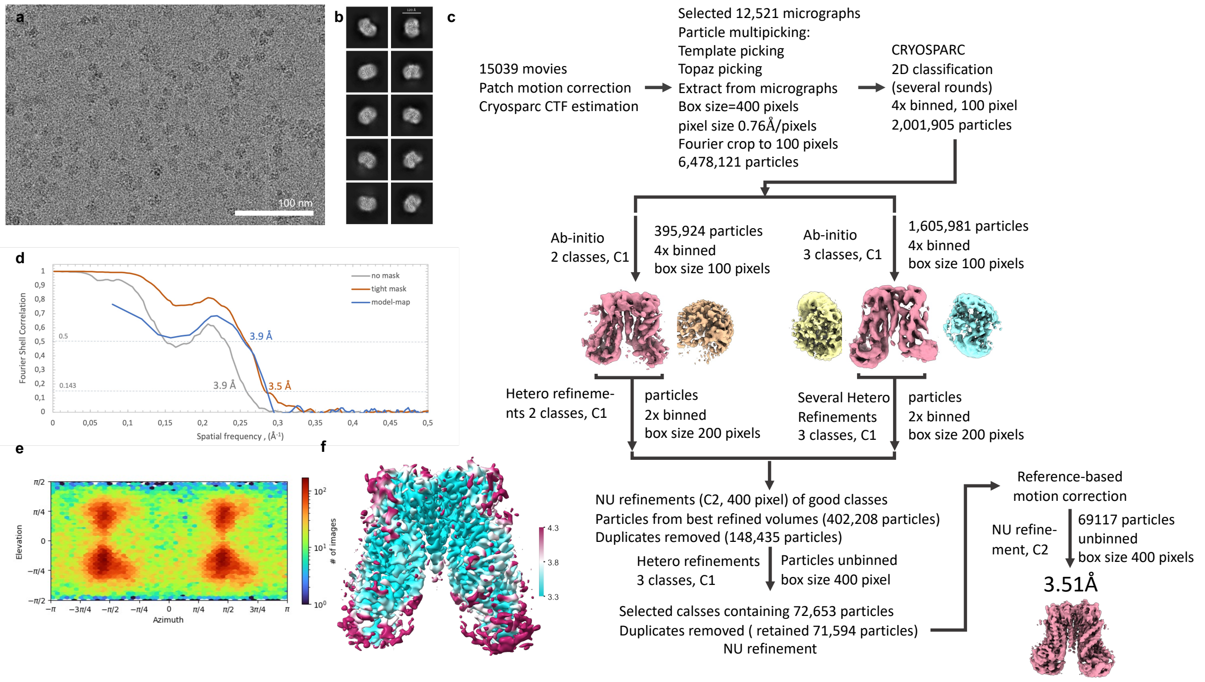


**Supplementary Figure 11 | Processing of TMEM16F-YFP purified and plunged using MISO. a**, A micrograph, **b**, Selected 2D class averages. **c**, Image processing scheme. **d**, Fourier Shell Correlation curves for unmasked and masked half-maps and between model and map. **e**, Heat map shows distribution of particle orientations. **f**, 3D map surface colored by local resolution.

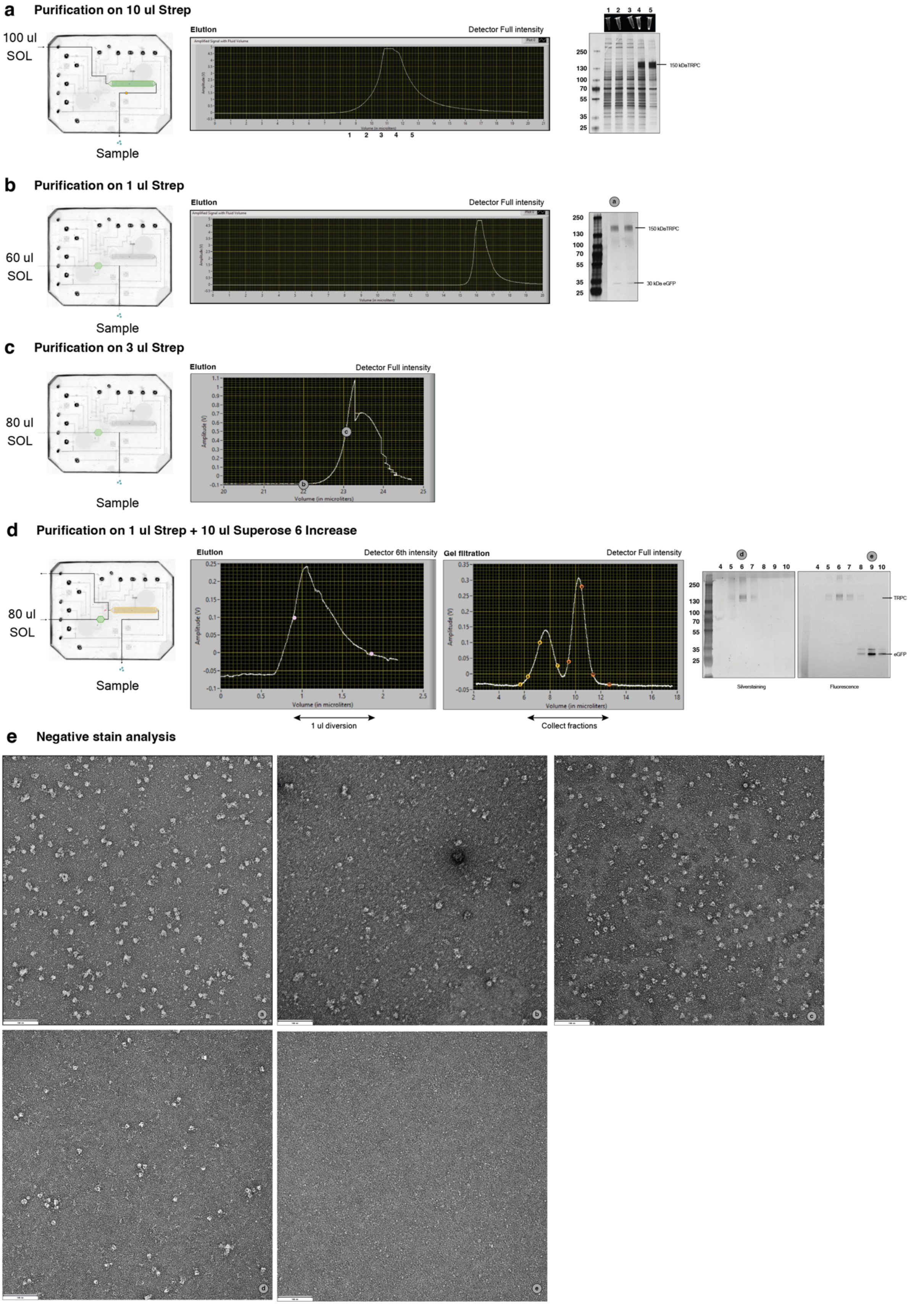


**Supplementary Figure 12 | Detailed overview of TRPC6 MISO purifications**. Chip designs used in the experiments are shown schematically, LabView fluorescent signal traces and SDS-PAGE analysis. **a,** 100 μl solubilizate (SOL) was loaded on 10 μl streptavidin resin and eluted with biotin. Fractions 1-5 analysed by SDS-

PAGE and visualised using silver stain. **b,** 60 μl SOL loaded on 1 μl strep column and eluted fractions were visualised using silver stain. **c,** 80 μl of SOL were loaded on 3 μl strep. Elution fractions were cryo-plunged. **d,** 80 μl of SOL were loaded on 1 μl strep. After elution, 1 μl of eluate was injected into the 10 μl Superose 6 Increase column. Collected fractions were analysed by SDS-PAGE visualised by fluorescence imaging and subsequently by silver stain. **e,** Fractions of the different steps during the optimisation were analysed by negative stain (a-e). The location of the fractions is shown on the LabView traces.

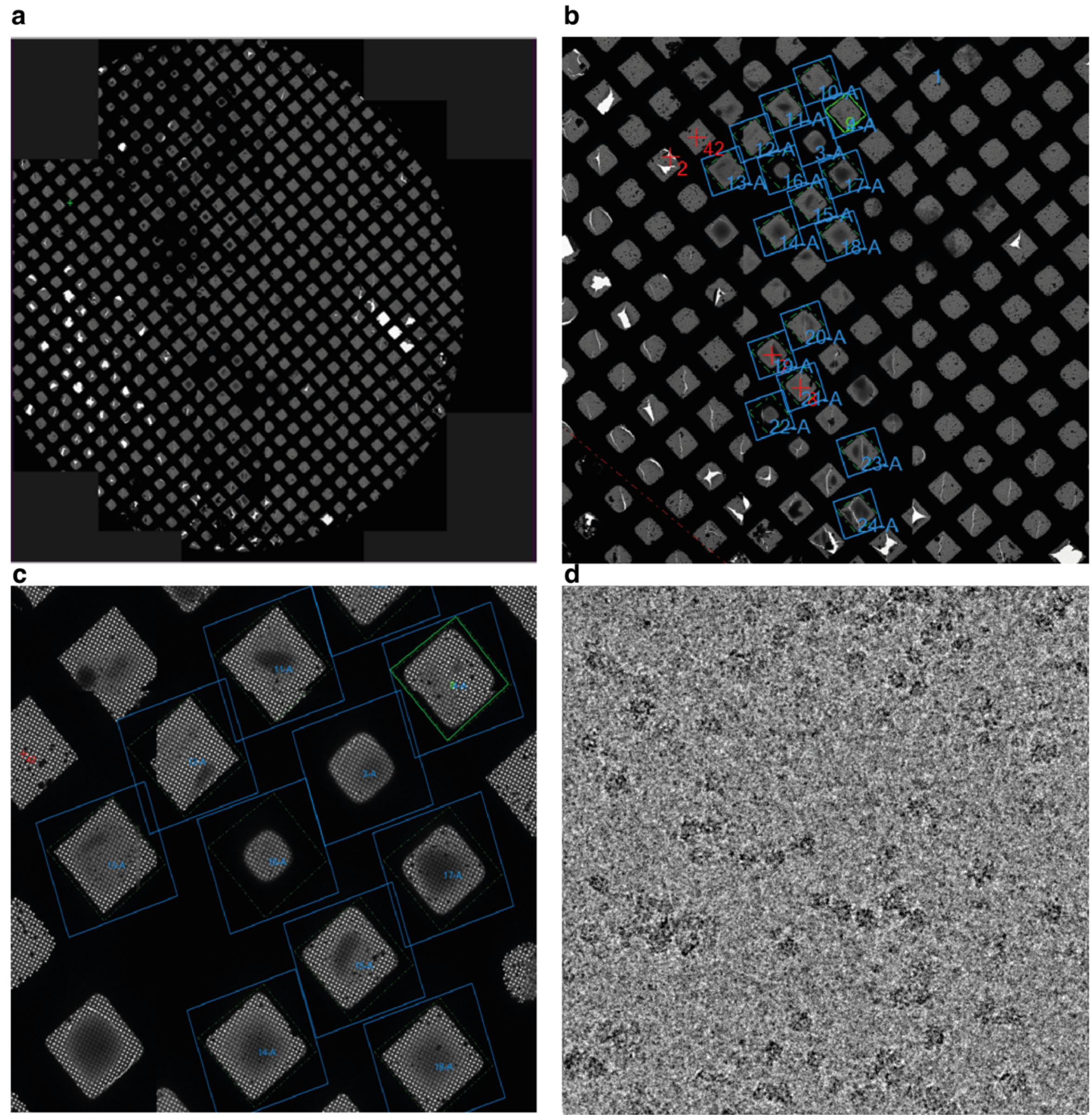


**Supplementary Figure 13 | Grid atlas and square montage of TRPC6 prepared using MISO. a,** Grid atlas of 80 nl sample deposited on gold UltraAUFoil R1.2/1.3 grid, automatically blotted for 0.5 sec with Whatman paper 4. **b, c**, Zoom in into the grid squares selected for cryo-EM data collection. **d**, An example of a micrograph from the dataset.

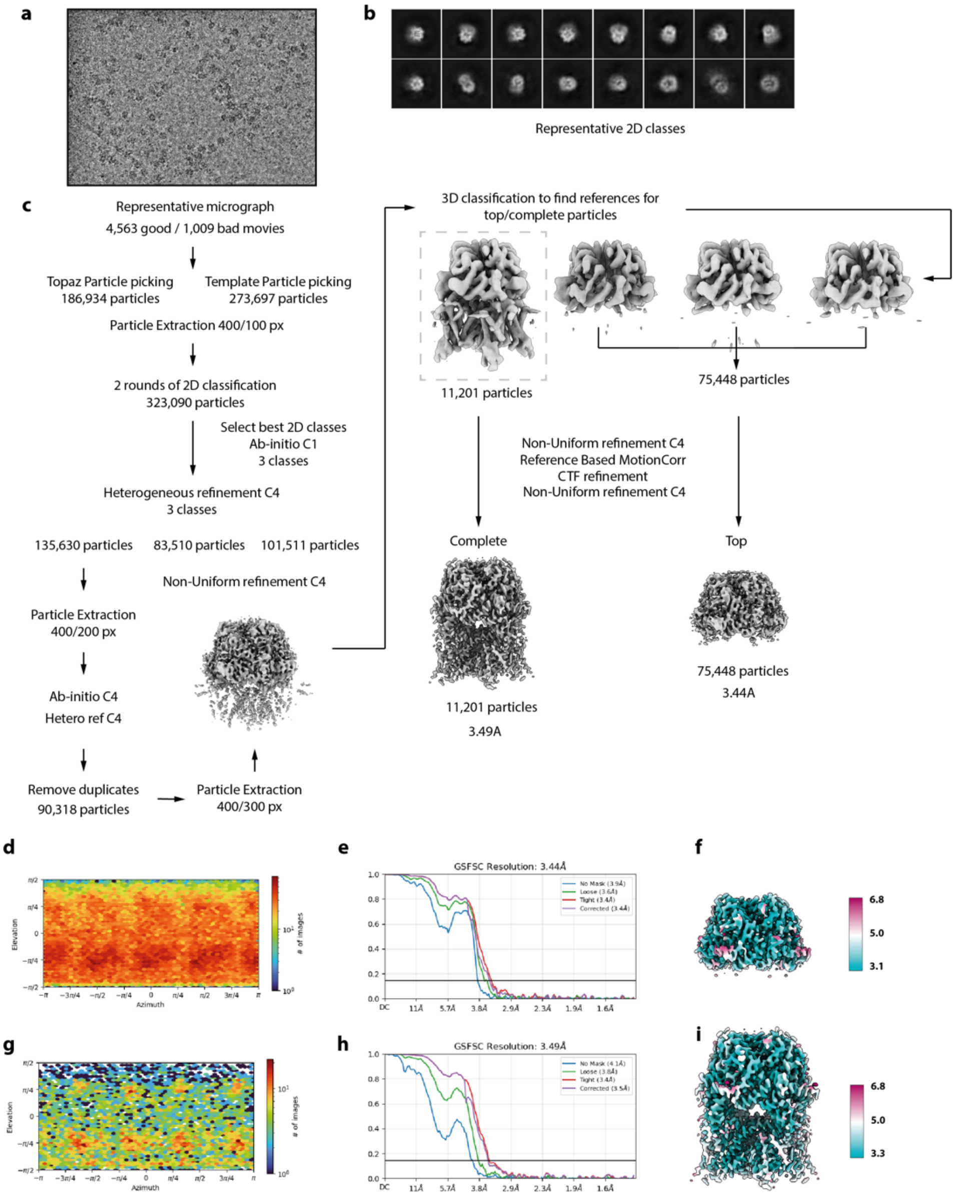


**Supplementary Figure 14 | Processing of TRPC6 prepared from 100 ul of starting material.** **a,** A cryo-EM micrograph and, **b** 2D class averages. **c**, Image processing scheme. **d, e**, Heat map shows distribution of particle orientations for soluble domain and complete trans-membrane complex, respectively. **e, h** Fourier Shell Correlation curves for unmasked and masked half-maps. **f**, **i** 3D map surface coloured by local resolution.

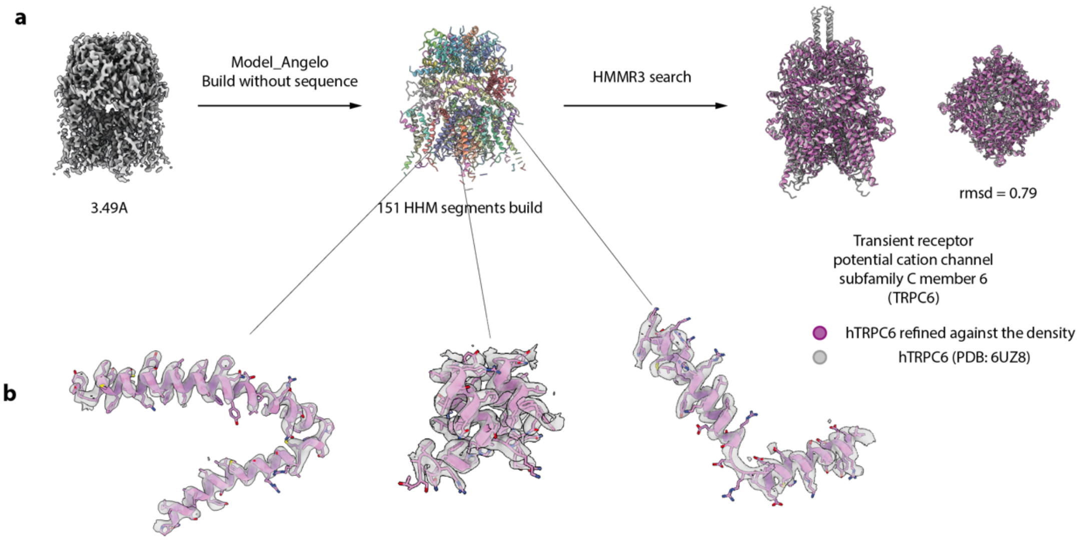


c

| v | E9PJN4 | Isoform of Q9Y210, Transient receptor potential cation channel subfamily C member 6 | | | | | | Homo sapiens | | | 7.6e-73 |
|---|---|---|---|---|---|---|---|---|---|---|---|
| Query | | Target Envelope | | Target Alignment | | Bias | Accuracy | % Identity (count) | % Similarity (count) | Bit Score | E-value |
| start | end | start | end | start | end | | | | | | Ind. / Cond. |
| 9 | 163 | 258 | 415 | 261 | 415 | 0.02 | 0.96 | 54.8 (85) | 81.3 (126) | 255.2 | 1.0e-72 / 6.0e-77 |

```
               .........*.........*.........*.........*.........*.........*.........*.........*
Query       9  ryekdsfefsksrinvykglaspafmtkgasnpvttsmeltekavlanldksyeedweenskqtrdfvtglldnvrnt. 87
               + ++dsf+ s+srin ykglaspa++   +++pv t++el+  lavlan+ k++++d+  +s q++dfv glld+ rnt
Target    261  KQKHDSFSHSRSRINAYKGLASPAYLSLSSEDPVMTALELSNELAVLANIEKEFKNDYKKLSMQCKDFVVGLLDLCRNTe 340
PP             4489**********************77888***************************66*******************6

               .........*.........*.........*.........*.........*.........*.........*......
Query      88  erveailigdseeipsdkdgrpnlerlyqalkydvkqfianpntkqqllsiwynnikglrnnseatyfliilalal 163
               e veail+gd e   s + grpnl+rl  a+ky+vk+f+a+pn+ qqllsiwy+n  glr+ ++a++fl++la a+
Target    341  E-VEAILNGDVETLQSGDHGRPNLSRLKLAIKYEVKKFVAHPNCQQQLLSIWYENLSGLRQQTMAVKFLVVLAVAI 415
PP             7.**************999******************************************************886
```

| v | Q6NV56 | Isoform of Q61143, Transient receptor potential cation channel, subfamily C, member 6 | | | | | | Mus musculus | | | 1.3e-36 |
|---|---|---|---|---|---|---|---|---|---|---|---|
| Query | | Target Envelope | | Target Alignment | | Bias | Accuracy | % Identity (count) | % Similarity (count) | Bit Score | E-value |
| start | end | start | end | start | end | | | | | | Ind. / Cond. |
| 6 | 100 | 97 | 193 | 98 | 191 | 0.00 | 0.96 | 53.2 (50) | 79.8 (75) | 137.5 | 1.9e-36 / 7.0e-41 |

```
               .........*.........*.........*.........*.........*.........*.........*.........*
Query       6  terfidsasrgnnpelreflkkekditinlsdrf.gfnalklsvangwlsiveillkksnlknvatalllslsqgyvriv 84
               +erf+d a +gn p +r++l +   + +n +  + g+nal+l+van +l+i e+llkk+nl  v+ alll++s+gyvriv
Target     98  EERFLDAAEYGNIPVVRKMLEECHSLNVN-CVDYmGQNALQLAVANEHLEITELLLKKENLSRVGDALLLAISKGYVRIV 176
PP             79**************************9.966************88**************77*****************

               .........*......
Query      85  tallkhpafasggnrl 100
               +a+l hpafa  g+rl
Target    177  EAILNHPAFAE-GKRL 191
PP             *********98.6*99
```

**Supplementary Figure 15 | Identification of the TRPC6 from cryo-EM map. a,** Model Angelo modelled sequence of 151 HHM segments into reconstructed cryo-EM map of the protein of interest. TRPC6 was identified as the highest likelihood target upon HHM profiles searched against the HMMER v3.3 reference proteome database. hTRPC6 was refined into the density (pink) and superimposed onto hTRPC6 (PDB 6UZ8, grey). **b**, Examples of density map regions with Michel Angelo sculptured fragments. **c**, Examples of HMMR search results for 2 sequences against the database.

**Supplementary Video 1 | Blotless deposition of protein solution on EM grid through capillary from MISO chip.**

**Supplementary Video 2 | MISO protein deposition on EM grid, blotting and plunging.**
